# Perturbomics of tumor-infiltrating NK cells

**DOI:** 10.1101/2023.03.14.532653

**Authors:** Lei Peng, Paul A. Renauer, Lupeng Ye, Luojia Yang, Jonathan J. Park, Ryan D. Chow, Yueqi Zhang, Qianqian Lin, Meizhu Bai, Angelica Sanchez, Yongzhan Zhang, Stanley Z. Lam, Sidi Chen

## Abstract

Natural killer (NK) cells are an innate immune cell type that serves at the first level of defense against pathogens and cancer. NK cells have clinical potential, however, multiple current limitations exist that naturally hinder the successful implementation of NK cell therapy against cancer, including their effector function, persistence, and tumor infiltration. To unbiasedly reveal the functional genetic landscape underlying critical NK cell characteristics against cancer, we perform perturbomics mapping of tumor infiltrating NK cells by joint *in vivo* AAV-CRISPR screens and single cell sequencing. We establish a strategy with AAV-SleepingBeauty(SB)- CRISPR screening leveraging a custom high-density sgRNA library targeting cell surface genes, and perform four independent *in vivo* tumor infiltration screens in mouse models of melanoma, breast cancer, pancreatic cancer, and glioblastoma. In parallel, we characterize single-cell transcriptomic landscapes of tumor-infiltrating NK cells, which identifies previously unexplored sub-populations of NK cells with distinct expression profiles, a shift from immature to mature NK (mNK) cells in the tumor microenvironment (TME), and decreased expression of mature marker genes in mNK cells. *CALHM2,* a calcium homeostasis modulator that emerges from both screen and single cell analyses, shows both *in vitro* and *in vivo* efficacy enhancement when perturbed in chimeric antigen receptor (CAR)-NK cells. Differential gene expression analysis reveals that *CALHM2* knockout reshapes cytokine production, cell adhesion, and signaling pathways in CAR-NKs. These data directly and systematically map out endogenous factors that naturally limit NK cell function in the TME to offer a broad range of cellular genetic checkpoints as candidates for future engineering to enhance NK cell-based immunotherapies.

## Introduction

Natural killer (NK) cells are lymphocytes with critical effector functions in innate immunity^1^ that do not require sensitization or specific antigens to initiate an effective immune response^2^. Effector populations of NK cells are able to lyse adjacent cells based on the expression of oncogenic transformation-associated surface markers^3^. In addition, regulatory NK populations can influence the functions of DCs^4, 5^, monocytes^6, 7^, T cells^8, 9^, and B cells^10, 11^ via cytokine production or through direct cell-cell contact in a receptor-ligand interaction-dependent manner^2, 12, 13^. These combined properties make the NK cell an attractive cell type of cancer immunotherapy^14^. The development of genetically engineered chimeric antigen receptor (CAR)-CAR have shown great therapeutic potential in NK cells^15, 16^, and compared with conventional CAR-T cell therapies, CAR-NK cells can use an allogeneic NK source without concern of GVHD and recognize tumor cells through cell native NK receptors, allowing for CAR-independent cancer elimination in tumors with antigen-loss^15, 17^. Currently (June 2022), there are over 35 clinical trials that involve testing of CAR-NKs against various hematological or solid tumor malignancies^18^. In several recent clinical trials, CAR-NK therapy has shown positive clinical trial outcomes against hematological malignancies^16, 19, 20^ and signs of promising potential for use in solid tumors^21, 22^.

Current forms of NK cell-based immunotherapy candidates face a number of obstacles, for example, the paucity^12^, lower proliferative capacity, and particularly decreased effectiveness, persistence or tumor infiltration^23, 24^. Various methods have been utilized to improve anti-tumor efficacy of NK cells, including *ex vivo* activation, expansion, and genetic modifications^25^. NK cells encode the same collection of ∼20,000 protein coding genes in their genome, many of which might play critical roles in regulating or limiting the anti-tumor function of NK cells. However, to date, there is no unbiased perturbation study in primary NK cells. Therefore, it is critical to create such a perturbation map of the NK genome, to guide the identification of new genes that can be targeted to enhance NK function, such as activation, proliferation, repression of inhibitory signals or exhaustion, persistence, or tumor infiltration.

Here, we performed a perturbomics study directly in primary NK cells, to systematically map thousands of genes for their quantitative effects in tumor infiltration, with a custom-designed high- density CRISPR library, targeting the surface proteome encoding genes embedded in an AAV-SB vector, in four different *in vivo* tumor models. Furthermore, we characterized the transcriptomic landscapes of tumor-infiltrating NK cells through single-cell RNA-sequencing (scRNA-seq) of tumor-infiltrating NK cells. Next, we leveraged an integrated analysis of the parallel functional genomics screens and the multi-parameter single-cell transcriptomic investigation of NK tumor-infiltration, which identified *Calhm2* as a convergent hit. Further *in vitro*, *in vivo,* and differential expression characterization showed that *CALHM2/Calhm2* knockout enhanced the anti-tumor function in both mouse primary NK and human CAR-NK cells.

## Results

### Multi-model, high-density *in vivo* perturbomics of tumor infiltrating NK cells

To systematically map the quantitative contribution of factors that influence NK cell tumor infiltration, we performed *in vivo* CRISPR-mediated knockout screens directly in primary NK cells using a custom high-density single guide RNA (sgRNA, gRNA) library, in four different tumor models (**Fig. 1a**). First, we designed Surf-v2, a high-density sgRNA library targeting the mouse homologs of the human surface proteome. Surf-v2 targets 2,863 genes with up to 20 sgRNAs per gene (56,911 gene-targeting sgRNAs). A total of 5,000 non-targeting control (NTC) sgRNAs reverse-ranked by potential off-targets from 500,000 random 20 nt sequences were spiked into the final library, for a total of 61,911 sgRNAs. We cloned Surf-v2 into the chimeric AAV-SB-CRISPR vector, which has high gene editing efficiency in primary immune cells. We then produce the AAV-SB-Surf-v2 viral library by packaging with AAV6 serotype system, and transduced primary Cas9-expressing splenic NK cells from constitutive transgenic Cas9 mice with a C57BL/6 (B6) background (**Fig. 1a**). These donor NK cells were intravenously (i.v.) injected into syngeneic host B6 mice pre-implanted with tumors. We performed these experiments with four tumor models in parallel, B16F10 melanoma, E0771 triple negative breast cancer (TNBC), GL261 glioblastoma and Pan02 pancreatic cancer (**Fig. 1a;** Methods). After 7 days of NK adoptive transfer, we extracted genomic DNA (gDNA) samples from pre-injection NK cells as well as tumors and spleens of tumor-bearing animals for screen readout (**Fig. 1a)**. Successful readout by next-generation sequencing (NGS) of the sgRNA library representations across all samples produced a dataset of multi-model, high-density *in vivo* perturbomics of tumor infiltrating NK cells (**Extended Data Fig. 1**).

**Figure 1:**
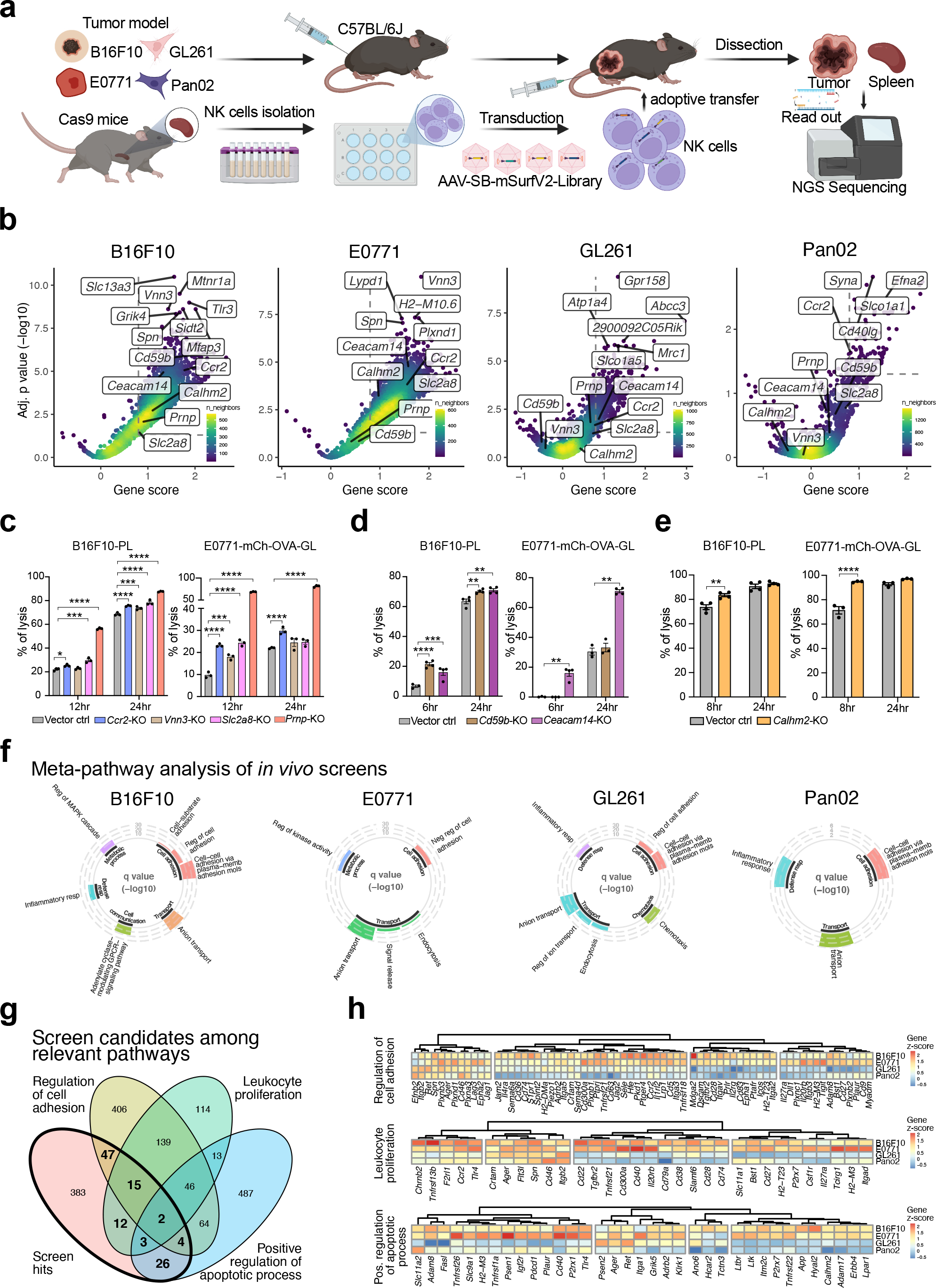
Functional genetic screens in four in vivo tumor models identifies candidate genes that enhance NK cell tumor infiltration. **a,** Schematic of the *in vivo* AAV-SB-Surf-v2 CRISPR KO screens for NK cell tumor infiltration performed in four independent syngeneic tumor models. **b,** Volcano plots of CRISPR-KO screen results for in vivo NK tumor-infiltration. Gene-level analysis results are shown as points with the FDR-adjusted p value and z-score represented by the position, and the result density presented by the color scale. **c-e**, Mouse primary NK cells with top hits knocking out showed better killing capability to cancer cells. Knockout *Ccr2*, *Vnn3*, *Slc2a8*, *Prnp* (**c**), *Ceacam14* (**d**), and *Calhm2* (**e**) in primary NK cells enhanced NK cells killing capability to B16F10-PL and E0771-mChe-GL cancer cells. Mouse primary NK cells with CD59b knockout (a top hit in B16F10 screen model but not E0771 screen model) showed enhanced killing to B16F10-PL cells but not to E0771-mChe-GL cells. **f,** Circular bar plots of meta-pathway analysis results for enriched genes of each *in vivo* NK screen. Meta-pathways are shown for relevant immune-related categories (bar color), and pathway significance is represented by bar height. **g,** Venn diagram of the overlap between screen result genes and select pathways. Gene hits were pooled across four CRISPR-KO screens and compared to three different gene ontologies that directly relate to tumor infiltration. **h,** Heatmaps of the CRISPR-KO screen enrichment (z-score) in genes of tumor infiltration-related pathways. Genes with positive z-scores in >2 in vivo screens are shown with bold font. Data are shown as mean ± s.e.m. plus individual data points in dot plots. Statistics: Mutiple t-test was used to assess statistical significance for comparisons; The p-values are indicated in the plots (**** p < 1e-4, *** p < 1e-3, ** p < 0.01, * p < 0.05). Source data and additional statistics for experiments are as a Source Data file.

With this dataset, we first performed a series of screen analyses. The overall library representation across the whole dataset showed sufficient retention of the full sgRNA library in the pre-injection primary NK cells, correlation between samples and models, as well as the dynamics of *in vivo* samples (**Extended Data Fig. 1a-c**). The tumor-model-specific sgRNA library representations revealed overall patterns of screen selection strength, as shown by the sample read distribution, cumulative distribution functions (CDFs) of samples groups, and principle component analyses (PCA) (**Extended Data Fig. 1c-e**).

We then analyzed these perturbation maps using a systematic approach, CRISPR-SAMBA (**Methods**), which incorporated generalized log-linear models (GLMs) and quasi-likelihood F-tests to assess which guides are enriched in the tumor, independent from the effects of the *in vivo* model (∼ intercept + *in vivo* samples + tumor samples, where *in vivo* samples = tumor or spleen samples) (**Extended Data Fig. 2a-c**). The sgRNA-level statistics demonstrated strong overall enrichment in tumor samples from the E0771 and B16F10 models, with moderate enrichment observed in GL261 and Pan02 models (**Extended Data Fig. 2d**). The sgRNA statistics were then aggregated into gene-level results, which showed enrichment in 1,131, 1,150, 175, and 32 genes in the B16F10, E0771, GL261, and Pan02 models, respectively (z-score > 0.8, q value < 0.01) (**Fig. 1b**). We also noticed greater similarity between the overall log-fold changes of the B16F10 and E0771, as well as between GL261 and Pan02 models (**Extended Data Fig. 3a**).

**Figure 2:**
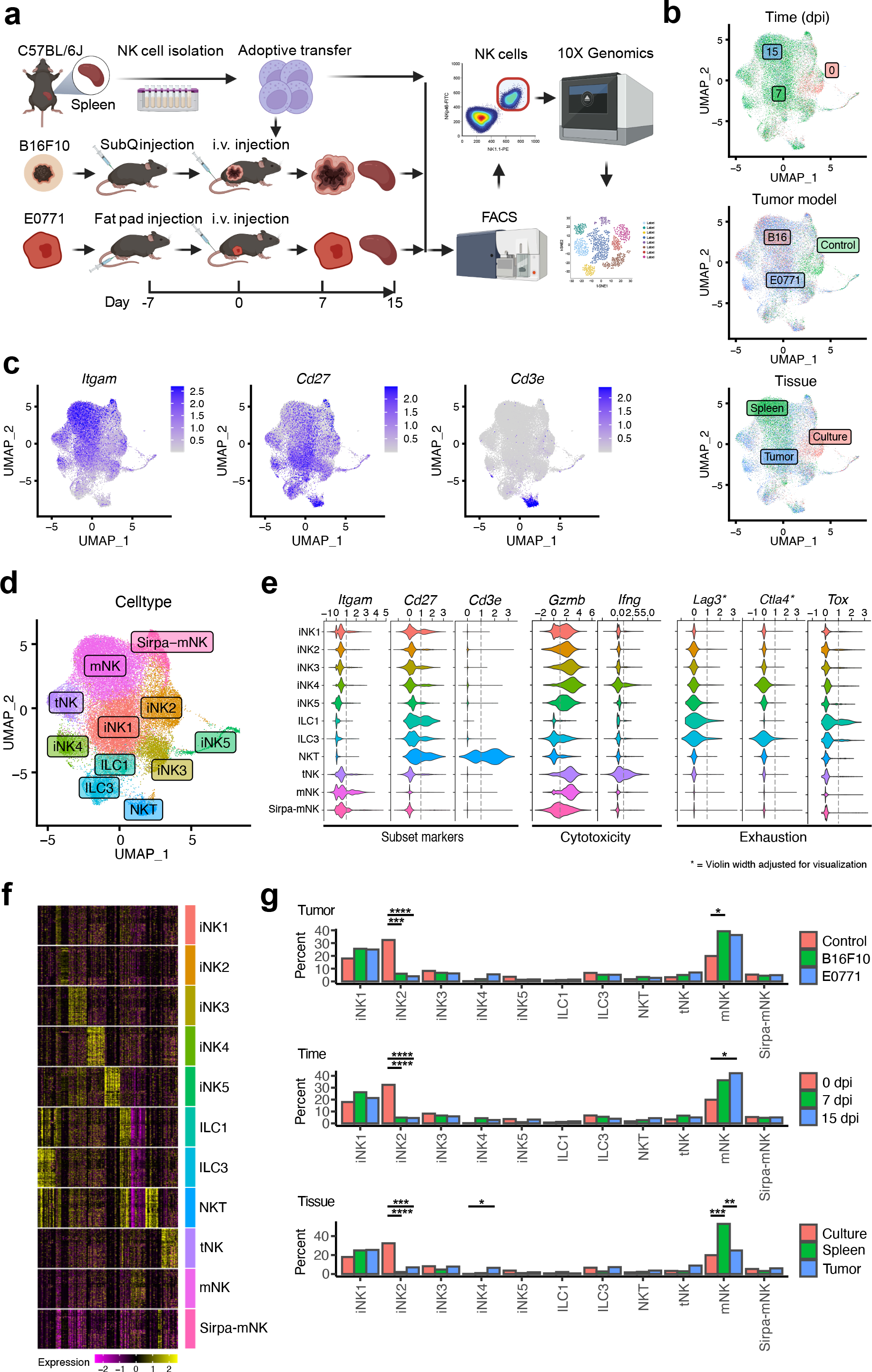
Single-cell transcriptomic analyses of in vivo NK cells across different timepoints, tissues and tumor models revealed unique NK population dynamics. **a,** Schematic for the single-cell transcriptomic exploration of NK cells within the tumor and spleen in two different in vivo cancer models, across multiple time points. **b,** UMAP plots of NK cells across nine integrated single-cell transcriptomic datasets. Cells are color-labeled according to their original dataset, including pre-transfer donor NKs (controls) and NK cells from different timepoints, tumor models, and tissues. **c,** UMAP plots of NK cells, color-coded by the single-cell expression of NK subset marker genes. **d,** UMAP plot of NK subset populations. **e,** Violin plots of the expression of select NK phenotype genes, compared across different NK subset populations. The distribution of log-scaled expression data is shown for each NK subset, and a dashed line represents a log-scale expression threshold of 1. **f,** Heatmap of discreet expression patterns across different NK cell population subsets. Scaled gene expression is presented for 100 representative cells of each NK population. **g,** Bar plots of NK subset percentages, compared across tumor-model, time, and the tissue type. Population comparisons were assessed by pairwise Chi-squared goodness-of-fit tests with FDR- correction (**** p < 1e-4, *** p < 1e-3, ** p < 0.01, * p < 0.05). Non-significant results were not labeled.

**Figure 3:**
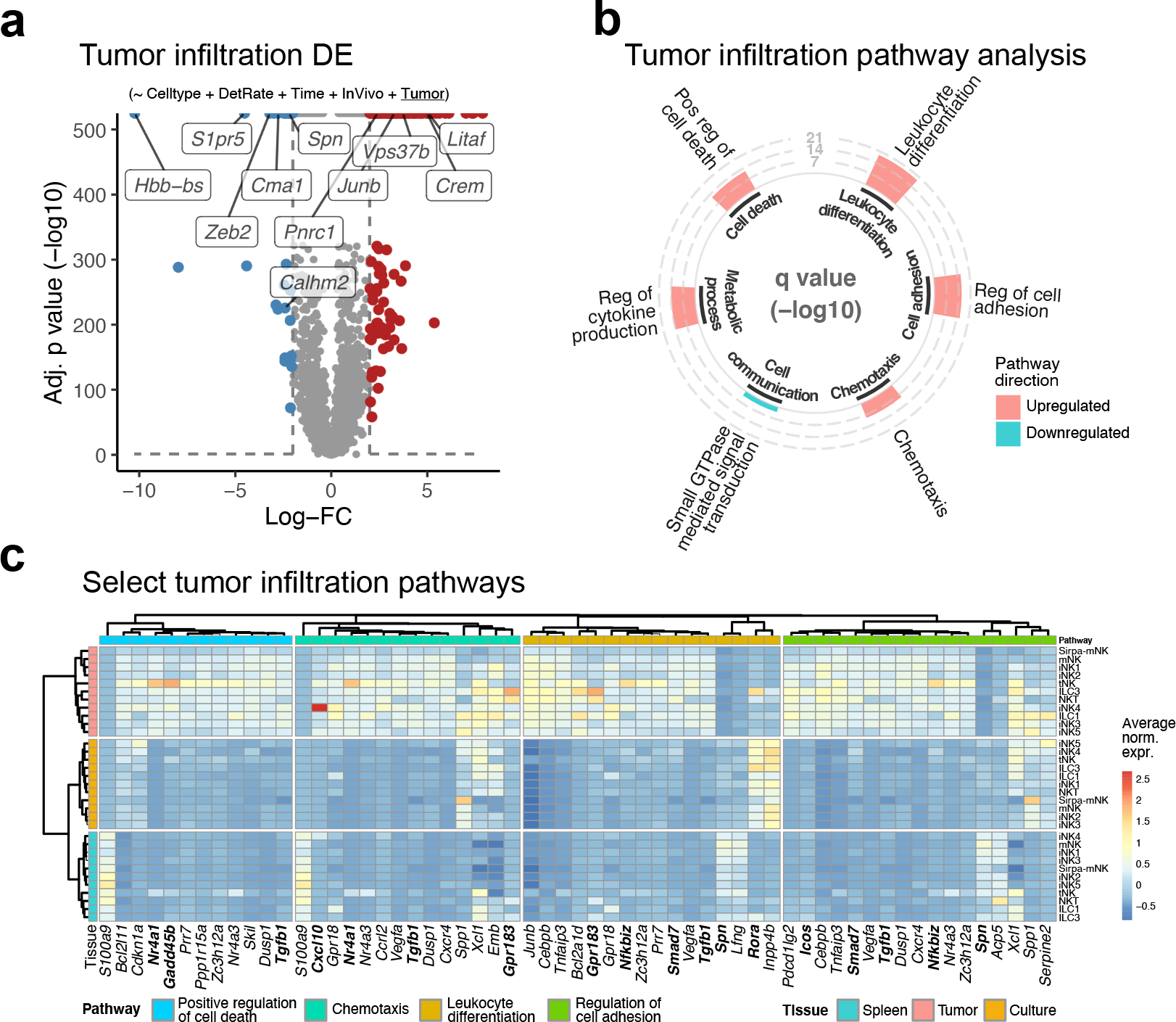
Single-cell DE analyses of tumor-infiltration revealed NK subset-specific pathway signatures **a,** Volcano plots of differential expression (DE) analyses for tumor infiltration of NK single-cell data. DE analyses were performed using single-cell expression data fitted to Gamma-Poisson generalized linear models (shown above plots), and quasi-likelihood F tests that assessed the effect of “tumor-infiltration” as a coefficient. Upregulated and downregulated genes are shown by respective red and blue dots (q < 0.01, absolute log-FC > 2), and the top 5 gene names are presented for each. **b,** Circular bar plots of meta-pathway analysis results for DE genes of the tumor infiltration analysis of NK single-cell expression data (top and bottom panels, respectively). Meta-pathways are shown for relevant immune-related categories, and pathway significance is represented by bar height. Meta-pathways from the analyses of upregulated and downregulated DE genes are indicated by red and blue bars, respectively. **c,** Heatmaps of select meta-pathways from the enrichment analysis of NK tumor infiltration DE genes (top and bottom panels, respectively). For each DE gene, population-averaged scaled- normalized gene expression is presented by color. Rows and columns are color-annotated by NK subset/tissue and meta-pathway, respectively. Hierarchical clustering was separately performed within each meta-pathway using highly variable DE genes (> 1 sd), and duplicate DE genes were allowed across meta-pathways.

We then assessed several screen hits by generating individual gene knockouts in primary Cas9+ NK cells, using the same AAV-SB-CRISPR vector expressing individual gene-targeting sgRNAs. We tested these individual gene knockout NK cells for their cancer lysis ability with *in vitro* co-culture assays using different cancer cell lines. Our results showed that deficiency of *Vnn3*, *Ccr2*, *Slc2a8*, *Prnp*, *Ceacam14*, and *Calhm2* (**Fig. 1c-e**) genes in NK cells, as compared to control NK cells, significantly enhanced their cytolysis of both B16F10-PL and E0771-mCh-OVA-GL cells. Knockout of *Cd59b*, enriched in the B16F10 cancer model alone, demonstrated greater cytolysis toward B16F10-PL cells but not E0771-mCh-OVA-GL cells (**Fig. 1d**).

Next, we used enriched screen results to determine relevant pathways of tumor infiltration across models. We analyzed gene enrichment using gProfiler2^26^, and aggregated enriched GO terms into meta-pathways (**Fig. 1f and Extended Data Fig. 4**). Pathway analysis results showed that anion transport was the only term common among the four models; However, there is also at least one meta-pathway for each model that relates to the regulation of cell adhesion. In addition, “cell-cell adhesion via plasma-membrane adhesion molecules” and “inflammatory response” meta- pathways are shared among B16F10, GL261, and Pan02 models.

**Figure 4:**
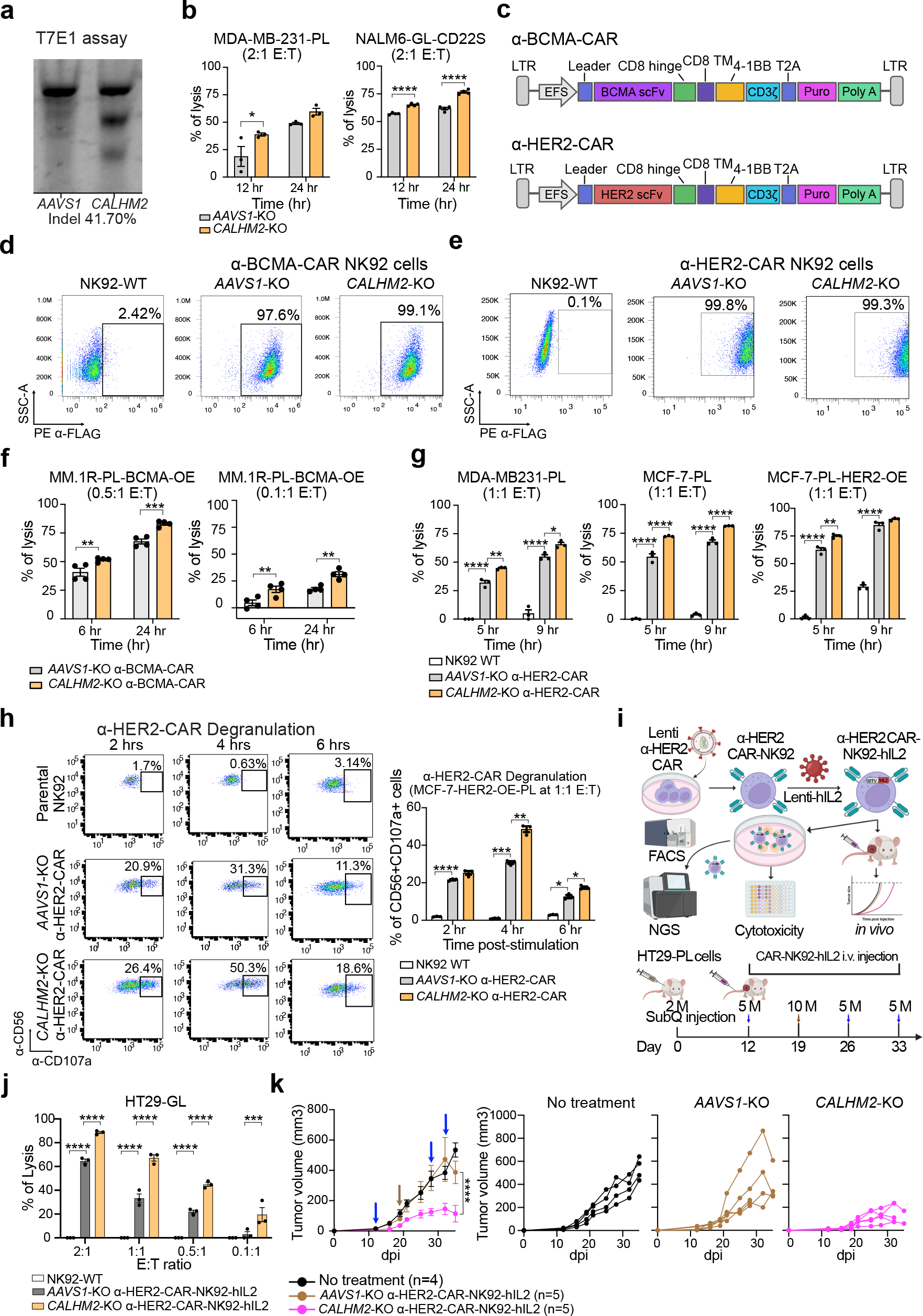
*CALHM2*-knockout in CAR-NK92 cells enhanced anti-tumor efficacy *in vitro* and ***in vivo*.** **a,** Gene editing of *CALHM2* in NK92 cells, as measured by T7E1 assay **b,** Enhanced killing capability of *CALHM2*-KO NK92 to MDA-MB-231 and NAML6-GL-CD22S cells. **c,** Illustration of α-BCMA and α-HER2 CAR construction. **d-e,** Flow cytometry plots of the percentages of *AAVS1*-KO and *CALHM2*-KO NK92 cells with (d) α-BCMA-CAR or (e) α-HER2-CAR expression. **f-g,** Enhanced killing capability of *CALHM2*-KO (f) α-BCMA-CAR-NK92 to MM.1R cells and (g) α-HER2-CAR-NK92 to MDA-MB-231, MCF-7-PL, and MCF-7-HER2-PL cells. **h,** *CALHM2*-KO α-HER2-CAR-NK92 cells secreted significantly higher CD107a when stimulated. **i,** (Top) Schematic of generation of α-HER2 CAR-NK92-hIL2 cells with *CALHM2* and *AAVS1* editing and functional validation with *in vitro* co-culture and *in vivo* tumor models. (Bottom) Timeline of *in vivo* tumor cytotoxicity analysis of *CALHM2*-KO α-HER2-CAR-NK92-hIL2 cells. **j,** Enhanced cytotoxicity of *CALHM2*-KO α-HER2-CAR-NK92-hIL2 cells to HT29-GL cells. **k,** (Left) Tumor growth curve of mice treated with *CALHM2*-KO or *AAVS1*-KO α-HER2-CAR- NK92-hIL2 cells. (Right) Spider curves of tumor growth for individual mice, plotted separated by group. In this figure: Data are shown as mean ± s.e.m. plus individual data points in dot plots. Statistics: Two-way ANOVA was used to assess statistical significance for multi-group curve comparisons; The p-values are indicated in the plots (**** p < 1e-4, *** p < 1e-3, ** p < 0.01, * p < 0.05). Source data and additional statistics for experiments are as a Source Data file.

Given the large number of hits from the four independent *in vivo* screens, we filtered the pooled list to include those with detectable expression in primary NK cells, based on ImmGen project data^27–30^. In addition, we further narrowed down gene sets that overlapped broad biological processes that might be relevant to tumor-infiltration by NK cells. These processes include (1) “cell adhesion”, as well as (2) “leukocyte proliferation” and (3) “positive regulation of apoptotic processes”, as a respective increase in NK cell expansion or survival could also result in increased numbers of NK cells within tumors (**Fig. 1g**). This filtering identified 109 genes, including several genes that serve as markers of exhaustion, such as *Tigit, Pdcd1/PD-1*, and *Lag3* ^31^ (**Fig. 1h**). Other notable hits include the Klrk1/NKG2D, an NK cell inhibitory receptor and marker of different NK cell developmental phases; CD27 of immature NK cells (iNK); as well as Itga1/Cd49a, Itga2/Cd49b, Itga3/Cd49c, and Spn/Cd43 of mature NK (mNK) (**Fig. 1h**). Taken together, this *in vivo* primary NK cell AAV-CRISPR screen dataset provided an overall quantitative perturbomics of surfaceome encoding genes in tumor infiltrating NK cells in four syngeneic tumor models, revealing a diverse collection of enriched hits as well as enriched pathways.

### Single-cell transcriptomic investigation of tumor-infiltrating NK cells

To further understand the tumor infiltration and behaviors of NK cell subsets, we performed single-cell RNA-seq analyses. Due to the strength of overall selection in the *in vivo* CRISPR screens above, we chose two tumor models, B16F10 and E0771 for single cell analysis of primary tumor infiltrating NK cells. Again, we established these orthotopic syngeneic tumor models by subcutaneous transplantation of B16F10 cells, and mammary fat pad transplantation of E0771 cells, into B6 mice. We then isolated donor splenic NK cells, also from B6 mice, without perturbations, and adoptively transferred them into tumor-bearing mice. After 7 days, we isolated NK cells from tumors and spleens at 7 and 15 days-post-injection by fluorescence assisted cell sorting (FACS), and subjected them to single-cell transcriptomics profiling using the 10X Genomics platform (**Fig. 2a**). Pre-transfer donor NK cells were also sequenced in parallel to serve as a control/baseline while exploring the effects of (1) time, (2) tumor type, and (3) tissue localization on NK cell phenotype (**Fig. 2b**).

We generated a total of nine different scRNA-seq datasets, represented by the various factors of time, tumor type, and tissue/localization (**Fig. 2a-b; Extended Data Fig. 5a**). We integrated all single cell data together based on dataset “anchors” that were identified using a reciprocal principal components analysis approach^32^. The integrated dataset was processed (**Methods**), and cell populations were visualized in reduced dimensional space using Uniform Manifold Approximation and Projection (**Extended Data Fig. 5a**)^33^. The cell populations were clustered by shared nearest neighbors (SNN) modularity optimization (Luvain algorithm with multilevel refinement) with the resolution optimized to ensure a unique transcriptional pattern between clusters, using highly variable genes (**Extended Data Fig. 5b-c**)^32, 34^. Cell populations were classified by the expression of known cell type-specific markers (**Extended Data Fig. 5c-d**), leading to the identification of various immune cell subtypes that were excluded from further analysis (**Extended Data Fig. 5c**). The remaining Ncr1+ cells were re-processed, visualized, and unbiasedly clustered (as previously described) (**Extended Data Fig. 6a**). Next, we broadly classified the NK sub-populations using relatively conservative definitions for murine NK cells, based on the expression of *Cd27*, *Itgam*, and *Cd3e* (**Fig. 2c and Extended Data Fig. 6b**). This resulted in the detection of 5 groups of immature NK cells (iNK; *Cd27*+*Itgam*-), 1 group of NK-T cells (*Cd3e*+), 1 group of transitional NK (tNK; *Cd27*+*Itgam*+), 2 groups of mature NK cells (mNK; *Cd27*-*Itgam*+), ILC1 (CD160+ Eomes+) and NCR+ ILC3 (Il7r+Kit+Rora+Gpr183+) (**Fig. 2d-e and Extended Data Fig. 6b**). It should also be noted, as *Itgam* was barely detectable in most cells, we considered tNK and mNK populations to be *Itgam*+, if more than 10% of the population had detectable expression. Unexpectedly, all of the iNK cells express some level of granzyme b gene (*GzmB*), while *Ifng* expression is only high in the tNK and iNK4 cells (**Fig. 2e and Extended Data Fig. 6b**), which is the only iNK subset with significantly increased *Cxcr4* expression, as well as no detectable expression of the *Spn* (*Cd43*) adhesion gene (**Extended Data Fig. 6c**). There are two ILC cell populations, each expressing *Lag3* and the *Tox* exhaustion TF, but *Tox* expression higher in ILC1 cells (**Fig. 2e and Extended Data Fig. 6b**). The ILC1 cells also express *Ctla4* and higher levels of *Pdcd1* (**Extended Data Fig. 5b**). Both exhausted populations express at similar levels of activating and inhibitory receptors, yet there are distinguishable differences in chemokine receptor expression, whereby ILC1 favors *Cxcr3* and ILC3 favors *Cxcr6* (**Extended Data Fig. 5c**). Single-cell analysis also revealed an mNK cell population that, uniquely among all other NK populations, expressed high levels of *Sirpa*, a gene recently identified as an NK immune checkpoint molecule^35^ (**Extended Data Fig. 6b**). The Sirpa-mNK cells are also marked by reduced detection of nearly all activating, inhibitory, chemokine, and adhesion receptors (**Extended Data Fig. 6c**).

**Figure 5:**
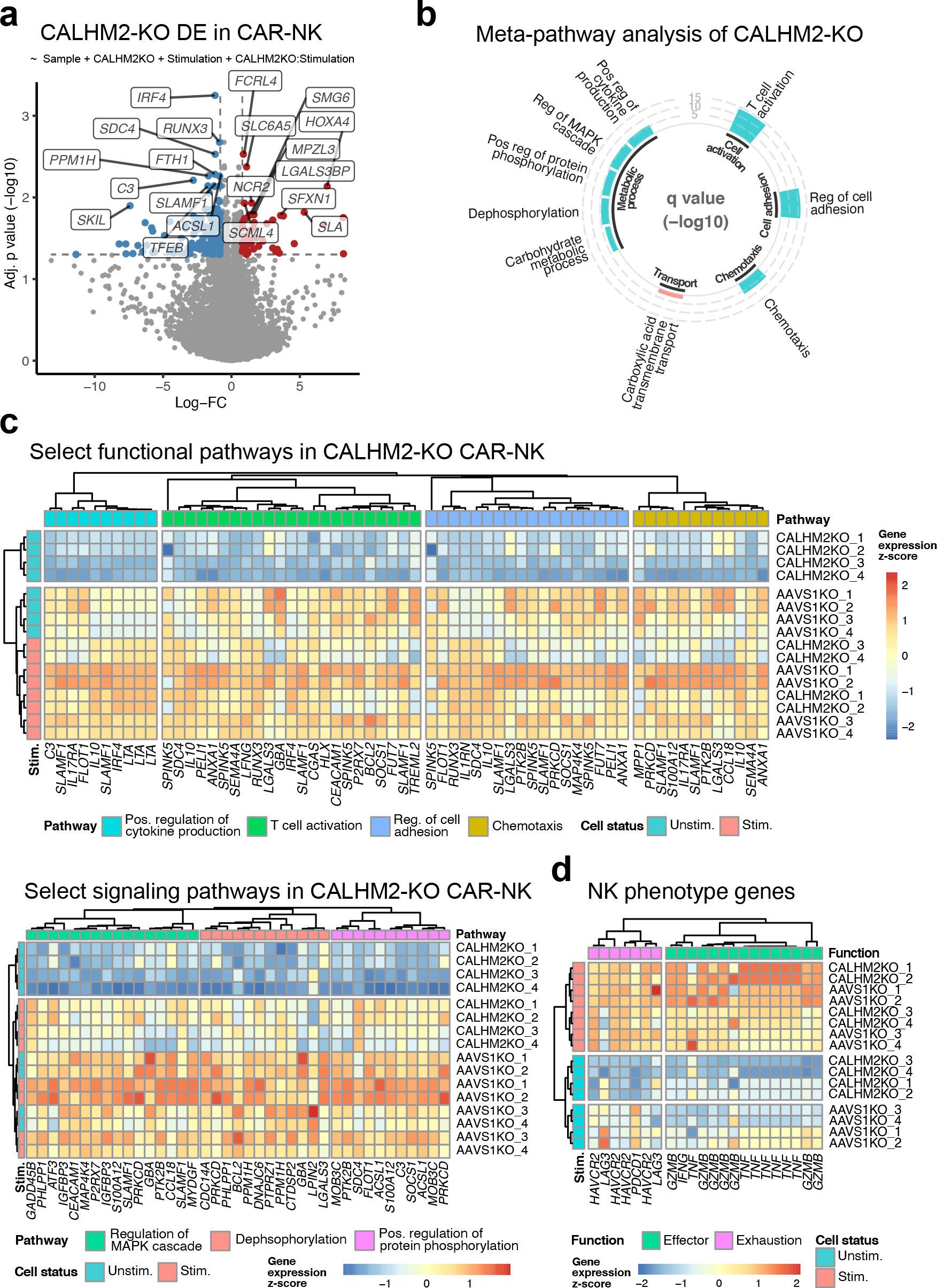
DE analysis in α-HER2 CAR-NK cells uncovered CALHM2-deficiency almost exclusively affects transcription in unstimulated cells **a,** Volcano plots of the differential expression (DE) analysis of CALHM2-deficiency in human donor α-HER2 CAR-NK cells. DE analysis was performed using bulk RNA-seq expression data fitted to a negative binomial generalized linear model that included coefficients for donor paired samples (CALHM2/AAVS1 gRNA), stimulation status, and the interaction between CALHM2- KO and stimulation. The effect of CALHM2-KO was assessed by quasi-likelihood F tests. Upregulated and downregulated transcripts are shown by respective red and blue dots (q < 0.05, absolute log-FC > 0.8), and the top significant DE gene names are labeled. **b,** Circular bar plots of meta-pathway analysis results for DE genes of the CALMH2-KO analysis in CAR-NK cell expression data. Meta-pathways are shown for relevant immune-related categories, and pathway significance is represented by bar height. Meta-pathways from the analyses of upregulated and downregulated DE genes are indicated by red and blue bars, respectively. **c,** Heatmaps of select meta-pathways from the enrichment analysis of CALMH2-KO in CAR-NK cells. The selected meta-pathways were grouped by functional pathways and signaling pathways (top and bottom panels, respectively). For each DE gene, scaled log-normalized gene expression is presented by color. Rows and columns are color-annotated by CAR-NK stimulation and meta- pathway, respectively. Hierarchical clustering was separately performed within each meta- pathway using highly variable DE genes (> 1 sd), and duplicate DE genes were allowed across meta-pathways.

### NK tumor population changes in progressing tumor models

NK cell population levels were quantified by single-cell transcriptomics across tumor progression at day 0, 7, and 15, in both melanoma and breast cancer models (B16F10 and E0771), and from different tissues. Our results indicated that the vast majority of NK cell subsets were highly stable across all conditions, allowing straight-forward detection of specific population shifts. In particular, our data showed that iNK4 cells (**Fig. 2g**) are also the only iNK subset specifically localized within the tumor (**Fig. 2g**). The iNK2 cell population also exhibited tumor-specific trends with its presence significantly associated with the pre-injection *in vitro* cell culture conditions, given the decreased presence in (1) each tumor model, (2) over time, and (3) in the spleen/tumor extracts (**Fig. 2g**). Each of these associations were observed along with an increasing abundance of mNK cells, except in tissue localization, where the significant loss of iNK2 is matched with an increase in only the splenic mNK (**Fig. 2g**).

### Expression patterns of tumor infiltration among NK populations

The transcriptional programs of tumor infiltration were explored in all NK cell populations by differential expression (DE) analyses using a Gamma-Poisson GLM to account for celltype, *in vivo* status, tumor infiltration, and the scaled RNA molecule detection rate of single cells. The DE analysis showed top upregulation in the early activation gene *Ly6a/Sca-1*^36^, and upregulation of the senescence-related *Litaf* gene^37^ (**Fig. 3a**). The top downregulated genes include mNK cell markers and genes involved in terminal NK differentiation, such as *Zeb2*, *Spn*, *S1pr5*, *Itgam*, *Prdm1*^38^ (**Fig. 3a**). Although there were significantly more mNK in the spleen than tumor and more intratumor iNK than mNK, this finding was still unexpected, given that the tumor-infiltrating NK population was comprised of ∼20% mNK (**Fig. 2g**), and the GLM should accounts for cell type. Therefore, we explored tumor-infiltration in only mNK cells populations via DE analysis and found a similar downregulation of terminally differentiated NK cell genes: *Zeb2*, *S1pr5*, *Spn*, *Ly6c2*, *Cx3cr1*, *Prdm1* genes (**Extended Data Fig. 7a**). The same trend was found in tumor-infiltrating iNK cells. The *Itgam* marker of the mNK subset was decreased in both iNK and mNK tumor-infiltrating cells, yet it was only significantly downregulated in the iNK. Among all tumor NK subtypes, there were also a consistent upregulation of the senescence-related *Litaf* gene^37^, and a consistent downregulation of *Calhm2*, a calcium-modulating enzyme gene^39^ that was also found as a hit in the *in vivo* AAV-CRISPR screens.

We then further investigated tumor-infiltration by NK cells with a meta-pathway analysis. The results showed that upregulated genes were significantly enriched in meta-pathways involved in cytokine production, leukocyte differentiation and the positive regulation of cell death, as well as tissue localization (chemotaxis and regulation of cell adhesion pathways) (**Fig. 3b**). Among these tumor-specific meta-pathways, there were consistent DE signatures of NK stimulation, such as *Junb*, *Cebpb*, and *Nr4a1/Nur77*, while there was a relatively consistent decrease in the meta-pathway expression signature of tumor Sirpa-mNK cells and increased expression signature in tNK cells (**Fig. 3c**). There were also clear NK population-specific differences across the meta- pathways, notably, a uniquely high expression of *Rora* and *Gpr183* in the ILC3. In addition, the tNK cells had distinctly higher expression *Nr4a1/Nur77*, strongly upregulated upon stimulation of NK activating receptors^40^. The iNK4 tumor cells also exhibited a distinct upregulation of *Cxcl10*, encoding a ligand of CXCR3 that has been shown to be upregulated in NK and NKT in a model of mycoplasma-enhanced colitis^41^.

Next, we investigated more specific behavioral differences among the major NK cell types. Meta-pathway analyses in both tumor iNK and mNK subsets showed positive enrichment of pathways for differentiation, inflammatory response, and the negative regulation of cell proliferation, while the mNK also had enrichment for the positive regulation of programmed cell death (**Extended Data Fig. 7b-d**). One contrast between the two NK classes was migration pathways, for which, tumor iNKs had positive enrichment for chemotaxis, whereas the mNKs showed negative regulation of leukocyte migration pathways. Taken together, these data suggest that the terminally differentiated mNK phenotype is negatively selected by the tumor microenvironment, even though the proportion of intra-tumor mNK cells increases over time (**Fig. 3f**)

### Single cell and gene expression signatures of tNK and iNK4 subsets with unique tumor functions

We performed DE analyses of tumor infiltrating tNK and iNK4, as each of these NK subsets seemed to have distinct expression signatures with exploring tumor NK cells as a whole. Although either DE analysis shared many of the top DE genes with tumor iNK/mNK, such as downregulated mNK markers (*S1pr5*, *Zeb2*, and *Spn*), and increased common activation genes (*Crem* and *Litaf*), there were unexpected changes in top upregulated genes, such as *Hspa1a/Hspa1b* heat shock protein genes in tNK (**Extended Data Fig. 8a**). The tumor iNK4 cells express high levels of the *Ctla4* checkpoint receptor, and *Spp1*, critical for long-lasting NK immune responses^42^, yet high expression is correlated with immunosuppression^43^ (**Extended Data Fig. 8a**). Meta-pathway analyses identified that tumor tNK cells had negative enrichment for activation and positive enrichment for the regulation of defense response, leukocyte differentiation, and the regulation of cytokine production (**Extended Data Fig. 8b-c**), for which tumor tNK had the highest expression of *Ifng* and *Ikbiz*, required for NK IFN-γ production in response to Il-12/Il-18 stimulation^44^. In addition, the splenic tNK still expressed *Ifng*, as well as high expression of *Egr3*, important for T and B cell activation, yet less is known for its role in NK cells^45^ (**Extended Data Fig. 8d**). In the tumor iNK4 cells, there is a positive enrichment for the inflammatory response differentiation, and regulation of adhesion and cell activation (**Extended Data Fig. 8b-c**). Within the meta-pathways, tumor iNK4 cells show a prominent upregulation of *Ifi204* and *Isg15* interferon response genes and of *Irf7*, the primary regulator of the type-I interferon response^46^ (**Extended Data Fig. 8d**).

### CALHM2 perturbation enhanced anti-tumor function in NK92 and CAR-NK92 cells *in vitro* **and *in vivo***

*CALHM2/Calhm2* scored both in the *in vivo* AAV-CRISPR screens and differential expression by single-cell transcriptomics in tumor infiltrating NK cells. Calcium homeostasis modulator protein 2, belongs to the calcium homeostasis modulator (CALHM) superfamily. *Calhm2* was found to regulate proinflammatory activity of microglial cells and is a potential therapeutic target for diseases related to microglia-mediated neuroinflammation^47, 48^; However, the function and anti-tumor roles of *CALHM2/Calhm2* in immune cells, especially NK cells, remains unclear. Our *in vitro* co-culture assay demonstrated enhanced cytotoxicity of *Calhm2-KO* mouse primary NK cells toward B16F10 and E0771 cancer cells (**Fig. 1e**). We then further characterized the effect of *CALHM2* deficiency in a clinically applicable human NK cell line: NK92^49^, which has been widely utilized for CAR-NK studies and has entered clinical trial stage. We first generated *CALHM2* mutant NK92 cells via Cas9/gRNA RNP electroporation and verified gene editing in the *CALHM2* locus by T7EI assay (**Fig. 4a**). NK92 cells with *CALHM2*-KO showed significantly increased killing of both MDA-MB-231 (breast cancer cell line) and NALM6-GL-CD22S cells (leukemia cancer cell line) in co-culture assays (**Fig. 4b**).

To further investigate whether *CALHM2* can serve as an endogenous gene target to enhance CAR-NK function, we established two different *CALHM2*-KO CAR-NK92 systems: α-BCMA-CAR and α-HER2-CAR separately (**Fig. 4c**) by lentiviral delivery. Puromycin selection was used to achieve near-complete (97.6% ∼ 99.8% purity) α-BCMA-CAR and α-HER2-CAR NK92 cells (**Fig. 4d-e**). Co-culture assays showed that *CALHM2*-KO α-BCMA-CAR-NK92 cells significantly kill more MM.1R-PL cells with BCMA overexpression (OE) at different effector : target cell (E:T) ratios (**Fig. 4f**), while *CALHM2*-KO α-HER2-CAR NK92 cells displayed a higher killing capability toward different breast cancer cell lines (MDA-MB231-PL, MCF-7-PL, and MCF-7-PL-HER2-OE) (**Fig. 4g**). In addition, degranulation assays showed that *CALHM2*-KO α- HER2-CAR NK92 cells expressed more CD107a after stimulation by cognate cancer cells (**Fig. 4h**). As NK92 cells are human interleukin 2 (hIL2)-dependent^50^, we established α-HER2-CAR- NK92-hIL2 cells via lentiviral delivery to assess *in vivo* efficacy (**Fig. 4i**). First, we confirmed that the addition of hIL2 does not alter enhanced cytotoxicity of CALHM2-KO by *in vitro* co-culture assays of *CALHM2*-KO α-HER2-CAR-NK92-hIL2 cells and HT29-GL cells (**Fig. 4j**). We then tested *CALHM2*-KO α-HER2-CAR-NK92-hIL2 cells’ therapeutic efficacy *in vivo* using a solid tumor model, induced by subcutaneous injection of an established human colon cancer cell line (HT29-GL) (**Fig. 4i**). Tumor growth kinetics showed that *CALHM2*-KO significantly enhanced *in vivo* anti-tumor efficacy of α-HER2-CAR-NK92-hIL2 cells, compared to *AAVS1*-KO controls, which showed limited to no efficacy compared with no treatment (**Fig. 4k**). Together, these *in vitro* and *in vivo* data demonstrated that *CALHM2* perturbation significantly enhanced the anti- cancer efficacy of human CAR-NK cells.

### CALHM2-knockout downregulates immune response pathways in unstimulated CAR-NK cells

To unbiasedly reveal how *CALHM2* perturbation changes CAR-NK function, we performed bulk mRNA-seq in *CALHM2-*KO (*CALHM2*-gRNA) and control (*AAVS1*-gRNA) human α-HER2 CAR-NK92-hIL2 cells, with and without stimulation (**Extended Data Fig. 9a-b**). The transcriptome patterns across the *CALHM2-*KO-CAR-NK dataset showed that the stimulated and unstimulated samples grouped separately by correlation analysis, while MDS visualization clustered samples according to stimulation and genotype (**Extended Data Fig. 9c-d**). We then identified the DE genes using the edgeR pipeline with a GLM for paired sample analysis along with *CALHM2*-KO, stimulation, and an interaction term between the two (**Fig. 5a and Extended Data Fig. 9c-d**), allowing for the effect of *CALHM2*-KO to be explored independent of stimulation. The *CALHM2*-KO effect in CAR-NK cells revealed 49 upregulated genes and 211 downregulated genes (FDR adjusted p < 0.05). Notable highly significant upregulated genes upon *CALHM2*-KO include *FCRL4*, an IgA-specific Fc receptor-like protein^51^, and NCR2 (NKp44), an NK activating receptor that senses platelet-derived growth factor (PDGF-DD isoform) from tumor cells^52^. Highly significant downregulated genes include *IRF4,* a key factor in exhaustion and differentiation in cytotoxic T cells^53^, and RUNX3, which regulates IL-15-dependent activation and differentiation in NK cells^54^.

Meta-pathway analysis of *CALHM2*-KO showed negative expression of chemotaxis, cell activation, as well as the regulation of MAPK, cell adhesion, and cytokine production pathways (**Fig. 5b**). Further inspection of the meta-pathway genes revealed an unexpected observation, whereby unstimulated *CALHM2*-KO samples had distinct downregulated expression patterns that separated these samples from all others via unsupervised hierarchical clustering (**Fig. 5c**). This downregulation was consistent among all relevant functional and signaling meta-pathways. Next, we assessed how the NK phenotype was affected by these transcriptional alterations, and identified that *CALHM2*-KO decreased overall effector and exhaustion molecule transcription in the unstimulated CAR-NK (**Fig. 5d)**; However, *CALHM2*-KO in the stimulated samples led to a slight decrease or negligible change on effector transcription, while it significantly decreased expression of *PDCD1/PD-1* and *HAVCR2/TIM3* exhaustion genes. DE analyses of the interaction term between *CALHM2*-KO and stimulation showed few significant genes, suggesting that the effect of *CALHM2*-KO and that of cancer stimulation in CAR-NK cells are largely orthogonal and independent of each other (**Extended Data Fig. 10a-b**). As expected, DE analyses of the cancer stimulation itself showed a substantial change of gene expression (**Extended Data Fig. 10a-b**). These data together revealed that *CALHM2-*knockout leads to a number of immune response pathways in CAR-NK cells at the baseline, in line with its effect in anti-tumor function.

## Discussion

NK-based cell therapy is a promising emerging branch of cancer immunotherapies. NK cell therapy leverages the advantages of rapid cytotoxic anti-tumor immune responses, TCR- independence, enhanced safety, simplicity in generating off-the-shelf allogeneic products, reduced off-target immune responses^55^, and reduced production of molecules associated with cytokine release syndrome (CRS) relative to other cell types^56–58^. Furthermore, the development of CAR-NK cells has increased the therapeutic potential of CAR-reprogramming by adding a reduced risk for alloreactivity and Graft-vs-Host Disease, potentially allowing for CAR-NK to be mass produced in a more cost-effective manner than CAR-T cells. Despite these promising attributes, NK cell-based immunotherapies still have many obstacles to overcome, including effective anti-tumor function, exhaustion, durable immune responses (persistence), and tumor infiltration. This requires rational engineering of substantially enhanced NK cells, particularly by modification of endogenous genes.

In order to systematically identify genes that can serve as endogenous targets to enhance NK function and thereby CAR-NK cell therapy, in this study, we functionally mapped tumor infiltrating NK cells with two independent, massive-scale investigations. First, we leveraged high-throughput, *in vivo* pooled AAV-SB-CRISPR knockout screens with a customized high-density sgRNA library, four separate *in vivo* tumor models, and functional genomics screen readout in tumor and spleen samples to quantitatively identify genes involved in NK tumor infiltration. The scope of these screens allowed our results to be robust across different cancer types, while the GLM-based analysis design enabled gene detection to be assessed independent of *in vivo* status.

Using this analysis method, our screens unveiled a perturbation map of thousands of surface protein encoding genes, and identified significant hits, including the benchmarking immune checkpoints and NK exhaustion markers (*Tigit*, *Lag3* and *Pdcd1*), along with a large collection of previously unknown / under-studied genes in NK cells.

Using single-cell transcriptomics, this study also performed an orthogonal, unbiased investigation of NK cell population structure and behaviors within the tumor microenvironment. We refined the scope, resolution, and applicability of our single-cell analysis by including two different tumor models, two timepoints, two tissues, and pre-transfer NK cells as a reference point in the analysis. These data have shown that there is a shift from iNK to mNK cells within the TME, despite decreased expression levels for key mNK marker genes in mNK cells. Our analyses identified previously unexplored sub-populations of NK cells, such as the iNK4 cells with distinct expression profiles of *Cxcr4* chemokine receptor and *Cxcr10*, which could have a supportive role in the control of tumors and infection through the recruitment of *Cxcr3*+ T cells and dendritic cells^41^. We also show that tumor-infiltrating NK cells have significantly increased regulation of cytokine production with genes related to TGF-β-signaling (*Tgfb1* and *Smad7*), which has a major immunosuppressive effect in NK cells^59, 60^; However, the specific molecules involved in the TGF- β-signaling and interactions with other signaling pathways is incompletely understood in NK cell populations^61^.

*CALHM2*/*Calhm2* emerged as both an enriched hit in the *in vivo* AAV-CRISPR infiltration screens, and as a consistently downregulated gene among tumor infiltrating NK cells from single cell sequencing. The role of CALHM2 in NK cells is relatively unclear, as it has primarily been studied in the context of Alzheimer’s Disease (AD)^39, 62–64^, and the known function of CALHM2 is context-specific, as the protein is a Ca2+-inhibited nonselective ion channel involved in calcium homeostasis and ATP release in depolarized cells^39^; However, a recent study in microglial cells began to reveal a new facet of CALHM2 function in immune cells, in which, its conditional knock- out decreased ATP-induced influx of calcium to the cytoplasm^48^. In turn, this inhibited activation of JNK, ERK1/2, NF-κB, and MAPK signaling, resulting in the decrease of IL-1β, TNFα, and IL- 6 production in AD but not WT microglia^48^. Given the strong relevance of these pathways in NK effector function, CALHM2 might has a similar role in unstimulated CAR-NK cells. This idea is supported by the consistent increase of effector transcripts in the control CAR-NK across condition and donor samples. CAR-NK cells also have tonic signaling, the continuous antigen-independent CAR signaling that promotes lymphocyte exhaustion^65^, which involves intracellular Ca2+ flux ^66^. Our functional and RNA-seq data revealed that (1) these transcription signals are downregulated in unstimulated CAR-NK upon CALHM2-deficiency, (2) CALHM2-deficiency has little effect on stimulated CAR-NK, and (3) CALHM2-KO CAR-NK cells had significantly decreased exhaustion.

## Methods

### Institutional Approval

This study has received institutional regulatory approval. All recombinant DNA and biosafety work were performed under the guidelines of Yale Environment, Health and Safety (EHS) Committee with an approved protocol (Chen-rDNA 15-45; 18-45; 21-45). All human sample work was performed under the guidelines of Yale University Institutional Review Board (IRB) with an approved protocol (HIC#2000020784). All animal work was performed under the guidelines of Yale University Institutional Animal Care and Use Committee (IACUC) with approved protocols (Chen 2018-20068; 2021-20068).

### Mouse models

The general health of the mice is in good condition (BAR: bright, alert and responsive) before the cancer-related experiments started. Mice were housed in a free access to water and food, temperature (approximately 22C) and humidity-controlled colony room, maintained on a 12h light/dark cycle (07:00 to 19:00 light on). Mice health checks were performed daily. Mice, both female and male, aged 8-12 weeks were used for experiments. Rosa26-Cas9-2A-EGFP constitutive expressed mice (Cas9 mice, also noted as Rsky/Cas9β mice) were used in this study. C57BL/6 and NOD-scid IL2Rgammanull (NSG) mice were purchased from JAX and bred in- house for *in vivo* tumor model and NK92 cell based therapeutic efficacy testing experiments.

### Cell lines

NK-92 cells were purchased from American Type Culture Collection (ATCC, Manassas, VA, USA). NK-92 cells and CAR-NK92 cells were cultured with MEM α, no nucleosides supplemented with 2 mM L-glutamine, 0.2 mM myo-inositol, 0.02 mM folic acid, 0.1 mM 2- mercaptoethanol, 200 IU/ml IL-2 (Biolegend), 12.5% FBS, 12.5% horse serum (Gibco, Life Technologies, America), and 1% penicillin/streptomycin. HEK 293T human embryonic kidney cells were bought from ATCC and cultured with DMEM medium containing 10% FBS and 1% penicillin/streptomycin. HT29, MCF-7, MDA.MB231 cells were transduced with a lentiviral virus (pXD023) that contains firefly luciferase and puromycin, yielding stable luciferase expressed HT29-PLuc, MCF-7-PLuc, MDA.MB231-PLuc cells. HT29-PLuc, MCF-7-PLuc and MDA.MB231-PLuc cells were cultured in DMEM medium with 10% FBS and 1% penicillin/streptomycin. NALM6 and MM.1R cells were culture in RPMI medium with 10% FBS and 1% penicillin/streptomycin and transduced with lentivirus (pXD024-contains GFP and luciferase) and lentivirus (pXD023-contains puromycin and luciferase) separately to generate NALM6-GL and MM.1R-PL cells. For NALM6-GL, MM.1R-PL, and MCF7-PL cell lines, cells were transduced with CD22-Blasticidin, BCMA-Blasticidin or HER2-Blasticidin lentivirus for overexpression of specific cancer antigen transgenes where appropriate, which were established by Blasticidin selection

### Mouse NK cells isolation and culture

Spleens were dissected from Cas9 or C57BL/6J mice, then placed into ice-cold PBS supplemented with 2 % FBS. Lymphocytes were released by grinding organs through a 100 mm filter, then re- suspended with 2 % FBS. Red blood cells (RBCs) were lysed with 1 mL ACK lysis buffer (Lonza) for 2 spleens at 1-2 min at room temperature, then neutralized with 2 % FBS PBS at 10 volumes per volume of lysis buffer. RBC-lysed lymphocyte solution was filtered through 40 mm filters to remove cell debris. NK cells purification was performed using EasySep™ Mouse NK Cell Isolation Kit (Stem Cell) according to the manufacturer’s protocols. NK cells were cultured at 0.5 x 10e6 cells / mL density in plates or dishes, with RPMI-1640 (Gibco) media supplemented with 10 % FBS, 2 mM L-Glutamine, 200 U / mL penicillin–streptomycin (Gibco), and 49 mM 2- mercaptoethanol (Sigma), hereafter referred to as cRPMI media, cRPMI media was supplemented with 50 ng / mL IL-2 (Biolegend), 50 ng / mL IL-15 (Biolegend).

### Design, synthesis and cloning of AAV-SB-Surf-v2 CRISPR library

A list of proteins in the human surface proteome was obtained from Bausch-Fluck et al^67^. The corresponding human genes were mapped to their mouse orthologous counterparts, for a total of 2867 genes. Exonic sequences for these mouse genes were obtained through Ensembl Biomart based on the mm10 genome assembly. Candidate Cas9 sgRNAs were then identified using FlashFry^68^, following default settings and using the scoring metrics “deonch2014ontarget”, “rank”, “minot”, “doench2016cfd”, and “dangerous”. With the resultant scoring matrix, sgRNAs were first filtered for those that did not have high GC content, no polyT tracts, and exactly one match in the mm10 genome. The sgRNAs targeting a given gene were then ranked by using the “doench2014ontarget” and “doench2016cfd” scores, by first converting each score to nonparametric ranks where high “doench2014ontarget” scores correspond to high ranks, while low “doench2016cfd” scores correspond to high ranks. The two nonparametric ranks were then added together, weighting the “doench2014ontarget” rank twice as heavily as the “doench2016cfd” rank. For final library design, all of the sgRNAs that are contained in the Brie library^69^ were first selected, then the composite ranks described above were used to choose the top scoring sgRNAs, up to a total of 20 sgRNAs per gene. The final set of on-target sgRNAs was composed of 56,911 sgRNAs targeting 2863 murine genes. A set of non-targeting control sgRNAs was designed by generating 500,000 random 20 nt sequences, followed by sgRNA scoring in FlashFry. The top 5000 non-targeting control sgRNAs were selected by choosing sgRNAs with a “doench2016cfd” score < 0.2 and < 100 total potential off-targets (maximum 4 mismatches). These 5000 control sgRNAs were added to the library, for a total of 61,911 sgRNAs. The oligo spacers for the surface-targeting gRNA library (Surf-v2) were generated by oligo array synthesis (CustomArray), PCR amplified, then oligos were cloned into double BbsI restriction digest sites of custom sgRNA vector by Gibson Assembly (NEB), after which, assembly products were transformed into high-efficiency competent cells (Endura) by electroporation (estimated library coverage = 233.6 fold). The custom sgRNA vector used in this study was a hybrid AAV-SB-CRISPR plasmid for targeting primary mouse NK cells (AAV-SB100x) that was constructed by gBlock fragments (IDT) followed by Gibson Assembly (NEB). The Surf-v2 library was cloned into the AAV-SB-CRISPR vector by pooled cloning to generate the AAV-SB-Surf-v2 plasmid library.

### AAV production

The AAV-SB-Surf-v2 library was packaged similarly to our previously described approach^70^. Low-passage HEK293FT cells were used for AAV production. Briefly, two hrs before transfection, D10 medium (DMEM (Gibco) medium supplemented with 10% FBS (Sigma) and 200 U/mL penicillin-streptomycin (Gibco)) was replaced by pre-warmed DMEM (FBS-free). For each 15cm-plate, HEK293FT cells were transiently transfected with 5.4 μg transfer, 8.7 μg serotype (AAV6) and 10.4 μg packaging (pDF6) plasmids using 130 μL PEI. After 6-12 h of transfection, DMEM was replaced with 20 mL pre-warmed D10 medium. Cells were dislodged and transferred to 50 mL Falcon tubes after 72 hr post-transfection. For AAV purification, 1/10 volume of pure chloroform was added and incubated at 37 °C with vigorously shake for 1 hr. NaCl was added to a final concentration of 1 M, shaking the mixture until all NaCl was dissolved, then pelleted at 20,000 x g at 4 °C for 15 min. The aqueous layer was gently transferred to another clean tube and discarded the chloroform layer. 10% (w/v) of PEG8000 (Promega) was added and shaken the tubes until dissolved. The mixture was incubated on the ice for 1 hr followed by centrifugation at 20,000 x g at 4 °C for 15 min. Supernatant was discarded and pellet was resuspended with 5-15 mL PBS including MgCl2 and benzonase (Sigma), incubated at 37 °C for at least 30 min. One volume of chloroform was added, shaken vigorously and spun down at 15,000 x g at 4 °C for 15 min. The aqueous layer was collected carefully and concentrated using AmiconUltra 100 kD ultracentrifugation units (Millipore). Virus was aliquoted and stored in -80 °C. To measure virus titer, RTqPCR was performed using Taqman assays (ThermoFisher) targeted to the human EFS promoter engineered in the AAV vector.

### AAV-SB-Surf-v2 NK cell *in vivo* screen in syngeneic tumor models

AAV-CRISPR screen was performed at >400x coverage. Naïve NK cells were isolated from the spleens of Cas9 mice. A total of 5e7 Cas9+ NK cells were transduced with AAV-Surf-v2 viral library. Syngeneic mouse models of melanoma, GBM, and colon cancer were set-up with subcutaneous injection of native B16F10, GL261, and Pan02 separately. Syngeneic mouse models of breast cancer were established by fat-pad injection of E0771 cells into C57BL/6J mice. AAV- Surf-v2 infected NK cells were adoptively cell transferred into tumor burden mice via i.v. (tail vein) injection. Four screen models were performed: B16F10 melanoma and E0771 breast cancer models, which reached an early endpoint (all mice euthanized by 20 days post tumor implantation (dpi), and GL261 GBM and Pan02 colon cancer models, which reached later endpoints (in GL261 model, all mice euthanized by 27 days post tumor implantation, and in Pan02 model, all mice were euthanized by 24 days post tumor inoculation).

### Tissue processing and genomic DNA extraction

Each mouse was dissected after euthanization. Spleen, dissected tumors, or pre-injected cell pellets were isolated for genomic DNA extraction. The genomic DNA extraction method follows our previously study^71^. Briefly, each sample was put in a 15 mL Falcon tube, 6 mL NK Lysis Buffer (50 mMTris, 50 mM EDTA, 1% SDS, pH adjusted to 8.0), and 30 μL of 20 mg/mL Proteinase K (Qiagen) were added to the tissue, and incubated at 55 °C overnight. After tissue disappeared, 30 μL of 10 mg/mL RNase A (Qiagen) was added to the lysed sample, and then tubes were inverted tubes 20 times and incubated at 37 °C for 30 min. Digested tissues were cooled on ice before adding 2 mL cold 7.5 M ammonium acetate (Sigma) to precipitate proteins. Samples were mixed thoroughly after adding ammonium acetate and vortexing for 10 s, followed by centrifuging at 4,000 x g at 4 °C for 15 min. After the spin, supernatant was removed to a new 15 mL Falcon tube and pellet discarded, 6 mL 100% isopropanol was added to the tube and inverted tubes until flocculent DNA was observed, centrifuged samples at 4,000 x g at 4 °C for 10 min. Genomic DNA pellets was washed one time with 70% ethanol, and then centrifuged at 4,000 x g at 4 °C for 5 min. The supernatant was discarded and removed remaining ethanol using a pipette. Air dry genomic DNA for 30-60 min, then added 0.5-1 mL nucleasefree H2O, resuspended DNA overnight at room temperature. The next day, gDNA solution was transferred to Eppendof tubes and measured concentration using a Nanodrop (Thermo Scientific). For cell pellets, 100-200 μL QuickExtract solution (Epicentre) was directly added to cells and incubated at 65 °C for 30 min. For mouse lymph nodes, QIAmp Fast DNA Tissue Kit (Qiagen) was used for gDNA extraction following the manufacturer’s protocol.

### AAV-SB-CRISPR NK screen readout and sequencing

Two rounds of PCR reactions were used for the sgRNA library readout. The first round PCR used genomic DNA (∼2 μg per reaction, three reactions per sample, ∼6 μg total per sample) to decrease the effect of PCR-amplification bias on the screen, and the second round PCR used 1 μL pooled PCR#1 product and barcoded primers. Each sample was amplified with different barcoded primers and pooled with equal quantity PCR products for Illumina sequencing. Primers for PCR#1^72^: Forward: 5’-aatggactatcatatgcttaccgtaacttgaaagtatttcg-3’ Reverse: 5’-ctttagtttgtatgtctgttgctattatgtctactattctttccc-3’ were used to amplify sgRNA cassette under cycling condition: 98 °C for 1 min, 25 cycles of (98 °C for 1s, 60 °C for 5 s, 72 °C for 12 s), and 72 °C 2 min for the final extension. All PCR reactions were performed using Phusion Flash High Fidelity Master Mix or DreamTaq Green DNA Polymerase (ThermoFisher). PCR#1 products for each biological sample were pooled and used for amplification with barcoded second PCR primers. The cycling condition of PCR #2 was: 98 °C for 1 min, 30-35 cycles of (98 °C for 1 s, 60 °C for 5 s, 72 °C for 12s), and 72 °C 2 min for the final extension. Second PCR products were pooled and then normalized for each biological sample before combining uniquely barcoded separate biological samples. The pooled product was then gel purified from a 2% E-gel EX (Life Technologies) using the QiaQuick Gel Extraction kit (Qiagen). The purified pooled library was then quantified with a gel-based method using the Low-Range Quantitative Ladder (Life Technologies), dsDNA High-Sensitivity Qubit (Life Technologies), BioAnalyzer (Agilent) and/or qPCR. Diluted libraries with 5-20% PhiX were sequenced with Illumina sequencers at the Yale Center for Genomic Analysis (YCGA).

### Single-cell RNA-sequencing of tumor-infiltrating NK cells

NK cells were isolated from the spleen C57BL/6J mice and expanded through *in vitro* culture. Syngeneic mouse models of melanoma and breast cancer model were set up with a subcutaneous injection of native B16F10 or a fat pad injection of E0771 cells into C57BL/6J mice. NK cells were adoptively transferred into tumor burden mice via i.v. (tail vein) injection. At 7 and 15 days after NK cells adoptively transferred, one mouse was sacrificed and the spleen and dissected tumor were collected. NKp46 and NK1.1 double positive NK cells were isolated by fluorescence-activated cell sorting (FACS), using FACS-Aria (BD). Sorted NK cells were counted and processed for single-cell RNA-sequencing library preparation by YCGA following manufacturer’s protocols.

### Lentivirus production

Lentivirus was produced using low-passage HEK239FT cells. One day before transfection, HEK293FT or HEK293T cells were seeded in 15 cm-dish at 50-60 % confluency. Two hrs before transfection, D10 media was replaced with 13 mL pre-warmed Opti-MEM medium (Invitrogen). For each plate, 450 μL of Opti-MEM was mixed with 20 μg CAR containing plasmid, 15 μg psPAX2 (Addgene), 10 μg pMD2.G (Addgene) and 100 μL lipofectamine 2000 (Thermo Fisher). After a brief vortex, the mixture was incubated for 15 min at room temperature and then added dropwise to the cells. After 6 hrs of transfection, Opti-MEM media was replaced with 20 mL pre-warmed D10 media. Viral supernatant was collected at 48 hrs post-transfection, then filtered using 0.45 μm filters (Fisher / VWR) to remove cell debris, and then concentrated using AmiconUltra 100 kD ultracentrifugation units (Millipore). All virus was aliquoted and stored in -80 °C.

### CRISPR gene editing in NK92 cells

The CRISPR-mediated knockout (KO) of *CALHM2* and *AAVS1* controls was performed by electroporation. Briefly, crRNA and tracrRNA were mixed in 1:1 ratio (final concentration 50 μM), heated at 95 °C for 5 min in a thermal cycler, then cooled to room temperature. 3 μL HiFi Cas9 protein (61 μM; Invitrogen) was mixed with 2 μL Buffer R for each reaction (Neon Transfection System Kit, Invitrogen), then mixed with 5 μL annealed crRNA:tracrRNA duplex, incubated the mixture at room temperature for 15 min. 3e6 of NK92 cells per reaction were resuspended in 100 μL Buffer R which included 10 μL RNP complex. 100 μL of cell:RNP mixture was loaded into the Neon Pipette without bubbles. The electroporation parameter was set at 1600 V, 10 ms for 3 pulses. Cells were immediately transferred to a 24-well plate with pre-warmed media after electroporation. KO efficiency for each target was examined after 5 days with T7E1 assay.

### CAR-NK92 cytotoxicity assay

To detect the cytotoxic capability of *CALHM2*-KO CAR-NK92 cells, cancer cell lines of NALM6- GL, MCF7-PL, MCF7-PL-HER2-OE, MDA-MB-231-PL, and MM.1R-PL were established as described above. The cancer cells were seeded in a 96-well plate first, then different Effector (NK92 cells): Target (cancer cells) ratio (E: T ratio) co-cultures were set up. Cytolysis was measured by adding 150 μg / mL D-Luciferin (PerkinElmer) using a multi-channel pipette. Luciferase intensity was measured by luminometer (PerkinElmer).

### CD107a degranulation assay

CAR-NK92 cells and NK92 cells were suspended with fresh culture medium supplied with 2 nM monensin and anti-CD107a-PE antibody (BioLegend) (1:1000 dilution), and stimulated with MCF-7-PL cells with E: T ratio of 1:1 (CARNK92: MCF-7-PL = 1:1) for 2 hrs, 4 hrs, and 6 hrs. At the end of co-culture, CARNK92 or NK92 cells were gently washed down with PBS and stained with anti-CD56-FITC for 30 min on ice, cells were analyzed using BD FACSAria.

### Bulk mRNA sequencing (mRNA-seq) library preparation

The mRNA library preparations were performed using a NEBNext® Ultra™ RNA Library Prep Kit, and samples were multiplexed using barcoded primers provided by NEBNext® Multiplex Oligos for Illumina® (Index Primers Set 2). *CALHM2*-KO anti-HER2 CAR-NK92-hIL2 cells and

*AAVS1*-KO anti-HER2-CAR-NK92-hIL2 cells were stimulated with HT29-PL cancer cells with 1:1 Effector (NK cells): Target (cancer cells) ratio. 4 hrs later, CAR-NK92 cells were sorted out and subsequently went for RNA extraction and mRNA-seq library preparations. Libraries were sequenced using a Novaseq 4000 (Illumina).

### *In vivo* animal experiments

NOD-scid IL2R-gamma-null (NSG) mice were purchased from JAX and bred in-house. Eight-to-twelve-week-old male mice were inoculated with 2e6 HT29-GL cells through subcutaneous injection. After 12 days, 5e6 *AAVS1*-KO or *CALHM2*-KO anti-HER2-CAR-NK92-hIL2 were injected intravenously into tumor burden mice. In the following days, CAR-NK92 cells were treated once a week and sequentially for 3 weeks. Treatment dose and time-point were labelled in the appropriate figures (**Fig. 4k**). Tumor progression was evaluated by tumor volume measurement by caliper, calculated as the following formula: vol = π/6 * length * width * height. All mice were sacrificed once they reached an endpoint according to the IACUC-approved protocols.

### CRISPR-KO screen data analyses

Raw sequencing data were demultiplexed and trimmed to the spacer sequences using Cutadapt v3.2^73^. The spacers were then aligned to the reference sgRNA library using Bowtie v1.3.0^74^, and aligned reads were compiled into a count matrix that was further processed in R using CRISPR- SAMBA^75^, which uses a modified pipeline of the edgeR differential expression analysis^76^ to calculate sgRNA enrichment, after which, sgRNA statistics are aggregated to acquire FDR-corrected p values and z-scores for each gene. For each tumor model, the CRISPR-SAMBA analysis was performed with tumor-infiltrating, splenic, and pre-injection control NK cell screen readout samples (default settings), using a quasi-likelihood (QL) generalized log-linear model (∼ InVivo + Tumor, where InVivo is any tumor or spleen sample) and a QL-F test with “Tumor” as the coefficient. For subsequent analyses, enriched genes were those with an FDR-adjusted p value < 0.05 and an absolute z-score > 0.8. In addition, the enriched genes were narrowed down to include those with detectable expression in NK cells, based on 18 NK samples from the ImmGen project (GEO: GSE122597)^27–30, 77^. ImmGen NK count data were processed by calculating the log2-transformed (0.5 pseudocount) gene-averaged counts-per-million, and log-expression > 1 was considered detectable expression.

### Single cell profiling

Splenocytes were collected from mice, and NK cells were purified by flow sorting, selecting NKp46+NK1.1+ cells with a FACSAria sorter. NK cells were then normalized to 1000 cells/μL. Standard volumes of cell suspension were loaded to achieve targeted cell recovery to 10000 cells. The samples were subjected to 14 cycles of cDNA amplification. Following this, gene expression (GEX), TCR-enriched and BCR-enriched libraries were prepared according to the manufacturer’s protocol (10x Genomics). All libraries were sequenced using a NovaSeq 6000 (Illumina) with 2x150 read length.

### Single cell transcriptomics data processing

Single-cell RNA-seq data were pre-processed with Cell Ranger v6.0.1 (10x Genomics) pipeline, using a standard pipeline that aligned reads to the mm10 mouse reference transcriptome and aggregated multiple datasets with the “agg” function. The aggregated datasets were subsequently processed using the Seurat v4.0.5 package for the R statistical programming language^34^. More specifically, each dataset was filtered to include cells with (1) 200-2500 RNA features, (2) < 5% mitochondrial RNA, (3) < 0.1% expression of Kcnq1ot1 (representing low-quality cells)^78^, and (4) < 5% combined expression of Gm26917 and Gm42418 (representing rRNA contamination)^79^. Each dataset was then log-normalized, scaled, and integrated via the Stuart et al. method, using the reciprocal-PCA dimensional reduction, 2000 anchors, and k = 20^80^. Integrated data were rescaled, and dimensional reduction was performed by uniform manifold approximation and projection (UMAP)^81^ using the first 27 dimensions from PCA, which were chosen by the inflection point of an elbow plot. Cells were clustered in low-dimensional space by generating a shared nearest neighbor (SNN) graph (k = 20, first 27 PCs) with modularity optimized using the Louvain algorithm with multilevel refinement algorithm (resolution = 0.2), based on the best spatial separation of major immune populations cells via *Cd3e*, *CD14*, *Cd19*, *Sdc1*, *Adgre1*, *Ncr1*, *Hbb-bs*, *Gypa, Pmel, H2-Aa, Ly6g, and Ptprc* expression (>10% of cell population expresses > 1 log-scale expression). NK cells were subset, rescaled, visualized by UMAP as before (first 20 PCs used), and clustered (resolution = 0.4), based on the separation of NK subset markers (*CD3e*, *Itgam*, and *CD27*) and exhaustion markers (*Lag3*, *Pdcd1*, and/or *Tox*) via UMAP and violin plot. These same markers were also used to label the cell clusters as specific NK populations using the same method as above. The labeled NK cell populations were assessed for within-cluster homogeneity by (a) performing Wilcoxon rank sum analyses of scaled expression data in each cluster compared to all other cells, (b) selecting the top 100 DE genes for each cluster (FDR-adjusted p value < 0.01, absolute log-FC > 1, cluster detection rate > 20%), and (c) determining the presence of discreet cluster-specific transcriptional patterns by hierarchical clustering and heatmap visualization.

### Single-cell differential expression analyses

Differential expression (DE) analyses of single-cell transcriptomics data were performed using a custom R pipeline, as previously described^82^. Briefly, raw single-cell data were filtered to include genes with detectable expression in >= 5% of cells, and then filtered data were fit to Gamma-Poisson generalized log-linear models (GLMs) (Source Data Fig. 3 and Source Data Extended Data Fig. 7-8) using the deconvolution method for the calculation of size factors^83, 84^. Differential expression analyses of fitted data were then assessed by empirical Bayes quasi-likelihood F (QLF) tests. GLM fitting and DE were performed using the glmGamPoi package for R^83^, assessing tumor-infiltration as the coefficients. For subsequent analyses, DE genes were those with an FDR-adjusted p value < 0.01 and an absolute log2 fold-change (log-FC) > 2.

### Bulk mRNA sequencing

Bulk mRNA sequencing was performed in HT29-stimulated and unstimulated αHER2-CAR-NK cells, in which there were paired AAVS1-KO and CALHM2-KO samples. Raw sequencing data were filtered and had adapters removed by Trimmomatic v0.39 in paired-end mode, clipping Illumina TruSeq adapters with the following settings: LEADING:3 TRAILING:3 SLIDINGWINDOW:4:20 MINLEN:30^85^. Trimmed, filtered reads were then aligned to the human transcriptome (GRCh38 Gencode version 96) using Kallisto v0.45.0 with the default settings^86^. Aligned reads were imported in R using the EdgeR package^76^, and the data were processed by removing low-expression transcripts with the filterByExpr command (default settings), normalized by trimmed mean of M-values method^87^. GLM dispersions were then calculated, and the data were fit to a quasi-likelihood (QL) negative binomial GLM for paired samples (∼ CALHM2-KO*Stimulation + Sample). Next, DE analyses were performed using empirical-Bayes QL F tests^76^, using either CALHM2-KO, Stimulation, or CALHM2-KO:Stimulation interaction as the coefficient. For subsequent analyses, DE genes were those with an FDR-adjusted p value < 0.05 and an absolute log2 fold-change (log-FC) > 0.80.

### Meta-pathway analyses

Meta-pathway analyses were performed using a modified pipeline of previously described strategy (Covid var paper). First, upregulated or downregulated DE genes were sorted by p-value and used as input for gene set enrichment analyses by the gProfiler2 R package with gene ontology (GO) terms for biological processes and known genes as the analysis domain^88, 89^. For enrichment analyses of the screen and bulk RNA-seq data, the threshold of DE was lowered to include more genes (screens: absolute z-score > 0.8, q < 0.05; bulk RNA-seq: absolute log-FC > 0.8, q < 0.01). Enrichment analysis results were filtered to keep significant GO terms (adjusted p value (gProfiler gSCS method) < 0.01), while excluding vague and poorly-matched GO terms (< 750 term genes, term overlaps >= 2 DE gene). If there were more than 2 filtered terms, analysis results were clustered into meta-pathways by generating an undirected network with (a) edges, weighted by similarity coefficients between genes of each term (coefficient = Jaccard + Overlap of genes between GO terms; coefficient threshold = 0.375), (b) a Fruchterman-Reingold layout, and (c) the terms were grouped by Leiden clustering (modularity optimization method, 500 iterations), using iGraph, network, and sna R packages. A representative “meta-pathway” was chosen from terms of each cluster, as the term with the highest precision value that was well-represented by the input gene list (term size >50 total genes, overlapping number of DE genes and terms is >10th percentile of filtered terms). The resolution for Leiden clustering was empirically optimized by the dataset type to limit the occurrence of redundant meta-pathways (resolution = 1.4 for bulk RNA-seq analyses, and 1.2 for the screen and scRNA-seq analyses).

For visualization, the five most significant meta-pathways were displayed as network plots with all clustered terms shown. In addition, circular bar plots were used to display functionally relevant meta-pathways, which (a) had at least one of the selected GO ancestor terms (leukocyte proliferation, cell activation, leukocyte differentiation, cell adhesion, chemotaxis, immune effector process, leukocyte migration, transport, cell communication, response to cytokine, defense response, metabolic process, regulation of apoptotic process, cell motility, cell death, and cell population proliferation) and were (b) filtered to include only the most significant of any meta- pathways with 100% DE gene overlap.

### Sample size determination

Sample size was determined according to the lab’s prior work or from published studies of similar scope within the appropriate fields.

### Replication

Number of biological replicates (usually n >= 3) are indicated in the figure legends. Key findings (non-NGS) were replicated in at least two independent experiments. NGS experiments were performed with biological replications as indicated in the manuscript.

### Randomization and blinding statements

Regular *in vitro* experiments were not randomized or blinded. Mouse experiments were randomized by using littermates, and blinded using generic cage barcodes and eartags where applicable. High-throughput experiments and analyses were blinded by barcoded metadata.

### Standard statistical analysis

Standard statistical analyses were performed using regular statistical methods. GraphPad Prism, Excel and R were used for all analyses. Different levels of statistical significance were accessed based on specific p values and type I error cutoffs (0.05, 0.01, 0.001, 0.0001). Further details of statistical tests were provided in figure legends and/or supplemental information.

### Data Collection summary

Flow cytometry data was collected by BD FACSAria.

All deep sequencing data were collected using Illumina Sequencers at Yale Center for Genome Analysis (YCGA).

Co-culture killing assay data were collected with PE Envision Plate Reader.

## Data analysis summary

Flow cytometry data were analyzed by FlowJo v.10.7.

All simple statistical analyses were done with Prism 9. All NGS analyses were performed using custom codes.

## Code availability

The code used for data analysis and the generation of figures related to this study are available from the corresponding author upon reasonable request.

## Data and resource availability

All of the generated data and analysis information/results for this this study are included in this article and its supplementary information files. In particular, source data and statistics for non- high-throughput experiments s are provided in Supplemental Tables. Processed data for NGS or omics data are provided in Supplemental Datasets. Raw sequencing data will be deposited to NIH Sequence Read Archive (SRA) or Gene Expression Omnibus (GEO). Original cell lines are available at commercial sources listed in supplementary information files. Other relevant information or data are available from the corresponding author upon reasonable request.

## Acknowledgments

We thank all members of the Chen laboratory, as well as our colleagues in Department of Genetics, Systems Biology Institute, Cancer Systems Biology Center, MCGD Program, Immunobiology Program, BBS Program, Yale Cancer Center, Yale Stem Cell Center, RNA Center and Center for Biomedical Data Sciences at Yale for assistance and/or discussion. We thank the Yale Center for Genome Analysis, Yale Center for Molecular Discovery, High Performance Computing Center, West Campus Analytical Chemistry Core, West Campus Mass Spec Core and Keck Biotechnology Resource Laboratory at Yale, for technical support. S.C. is supported by NIH/NCI/NIDA (DP2CA238295, R01CA231112, U54CA209992-8697, R33CA225498, RF1DA048811), DoD (W81XWH-17-1-0235, W81XWH-20-1-0072, W81XWH-21-1-0514), Damon Runyon Dale Frey Award (DFS-13-15), Melanoma Research Alliance (412806, 16-003524), St-Baldrick’s Foundation (426685), Breast Cancer Alliance, Cancer Research Institute (CLIP), AACR (17-20-01-CHEN), The Mary Kay Foundation (017-81), The V Foundation (V2017-022), Alliance for Cancer Gene Therapy, Sontag Foundation (DSA), Pershing Square Sohn Cancer Research Alliance, Dexter Lu, Ludwig Family Foundation, Blavatnik Family Foundation, and Chenevert Family Foundation. PR is supported by Yale PhD training grant from NIH (T32GM007499) and Lo Fellowship of Excellence of Stem Cell Research. JJP is supported by NIH MSTP training grant (T32GM007205). RC is supported by NIH MSTP training grant (T32GM007205) and NRSA fellowship (F30CA250249).

## Author Contributions

SC conceived the study and designed with LP, PAR and LpY. LP and LpY performed most of experiments. PAR performed most NGS analysis. LjY, YZ, QL, MB, AS, SZL assisted experiments. JJP, YZ performed certain NGS analysis. RDC designed Surf-v2 library. LP, PAR, and SC prepared the manuscript with inputs from all authors. SC secured funding and supervised the work.

## Extended Data

**Extended Data Figure 1:**
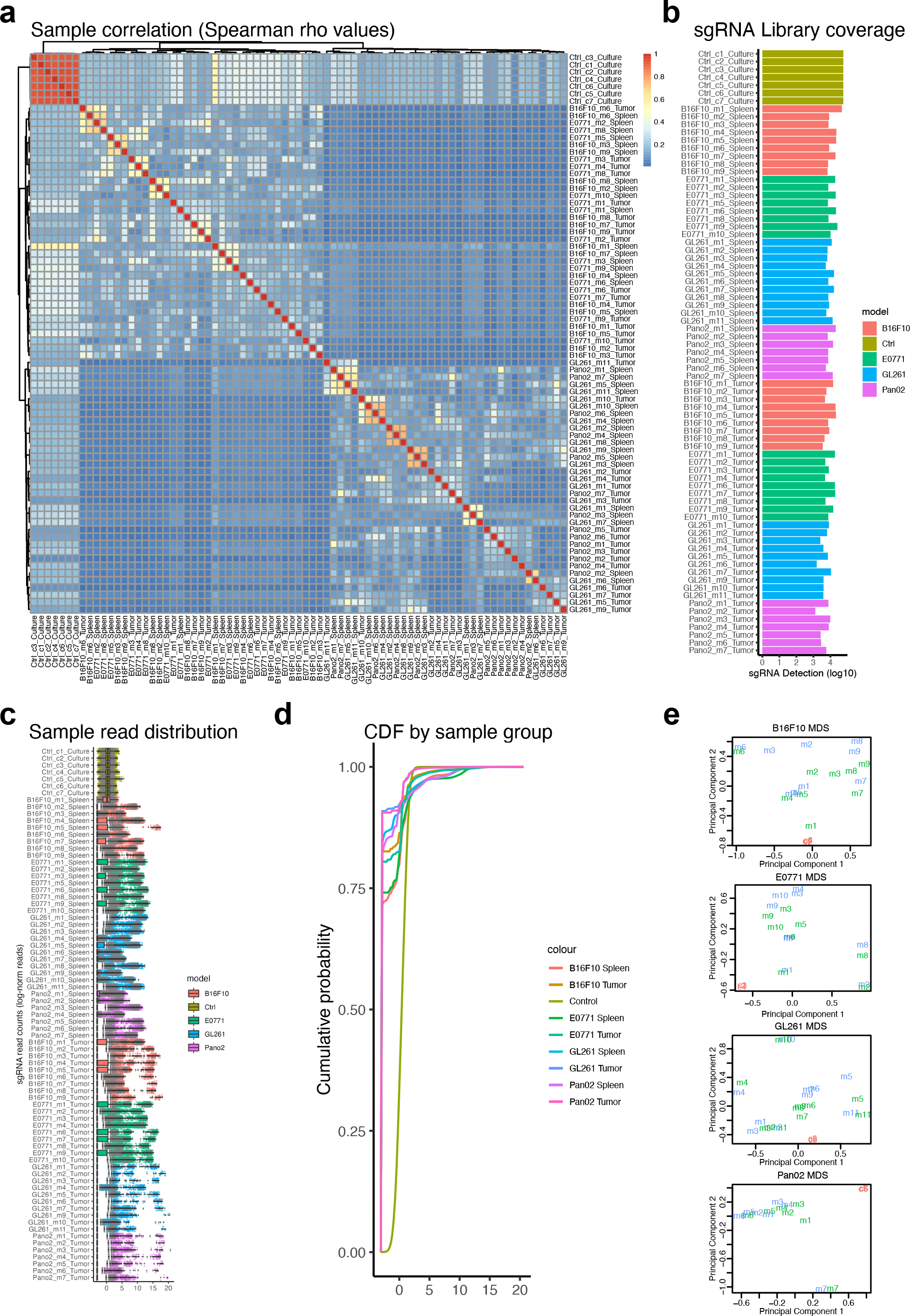
Quality control metrics of four independent in vivo CRISPR-KO screens for NK cell tumor infiltration. **a,** Heatmap of the Spearman correlation between samples of the in vivo CRISPR-KO screens for NK cell tumor infiltration performed in four independent tumor models. **b,** Bar plot of the sgRNA library coverage for all screen samples. sgRNA detection is represented by bar height, while tumor models are depicted by bar color. **c,** Box-whisker plot of sgRNA read count distributions across all screen samples. The tumor model is depicted by box and dot color. Non-targeting control (NTC) sgRNAs are shown as grey dots. **d,** Line graphs of the cumulative distribution function of screen samples, pooled by tumor model and the tissue from which the NK cells were extracted. **e,** Scatter plots for the multidimensional scaling of processed screen data of each tumor model.

**Extended Data Figure 2:**
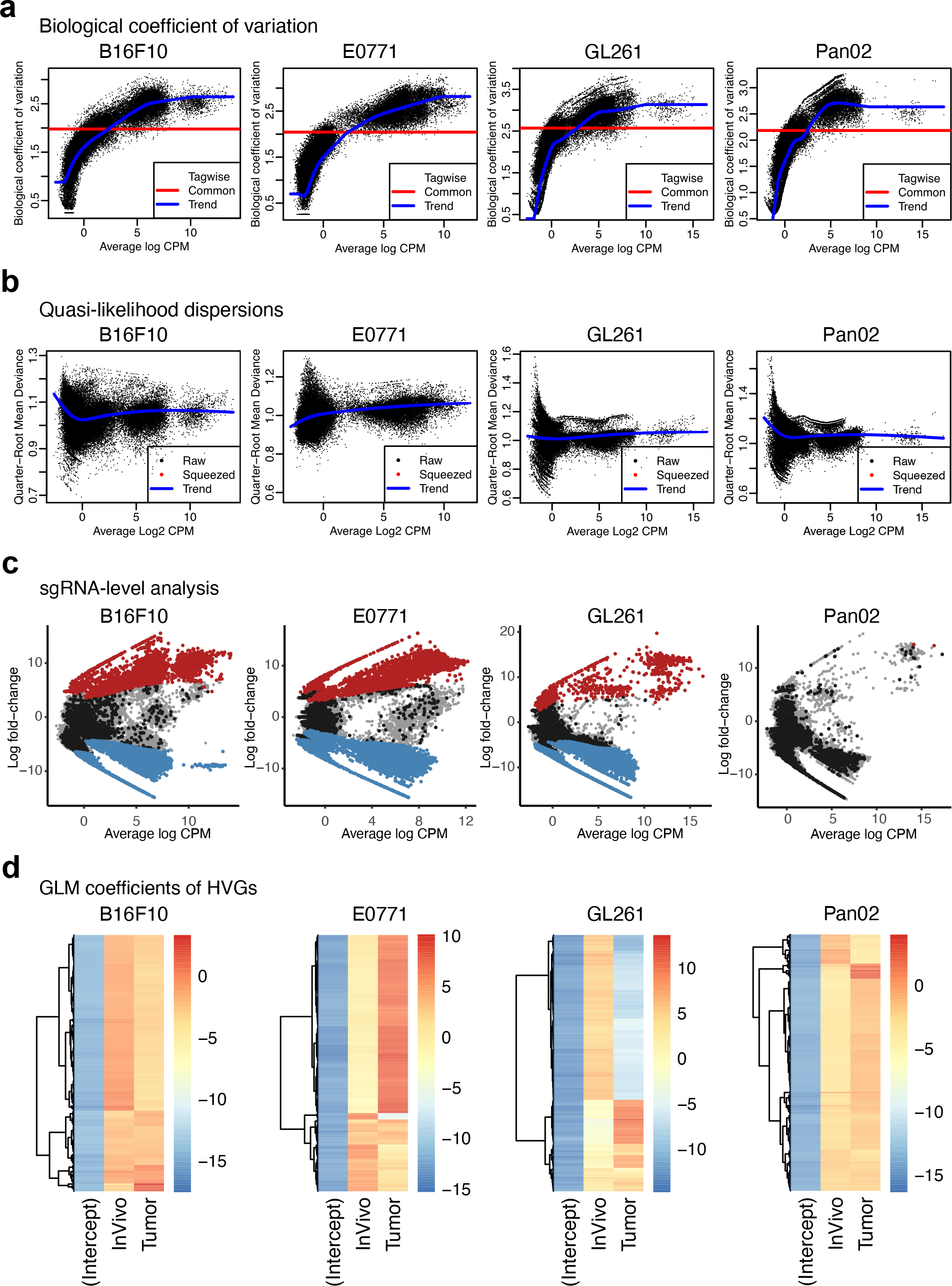
Quality control metrics for the CRISPR-SAMBA analyses of four independent in vivo CRISPR-KO screens for NK cell tumor infiltration. **a,** Scatter plots of the dispersion estimates for the biological coefficient of variation (BCV) in the *in vivo* CRISPR-KO screen analyses for NK cell tumor infiltration. The BCV is calculated as the square root of the negative binomial dispersion, and the common, trended and gene-wise BCV are presented relative to the log2 read counts per million (CPM). **b,** Scatter plots of the quasi-likelihood (squeezed) dispersions used for screen analyses in different tumor models. The quarter-root mean deviances of the raw and squeezed negative binomial dispersions, along with the trended dispersion line, are presented relative to average sgRNA read count abundance (log2 CPM). **c,** MA plots of the sgRNA-level CRISPR-KO screen analyses for NK cell tumor infiltration in four tumor models. The log fold-changes of each sgRNA are depicted relative to the read count abundance (log2 CPM). Non-targeting control, significantly enriched, and significantly depleted sgRNAs are visualized as black, red, and blue points, respectively. **d,** Heatmaps of the experimental coefficients in the generalized log-linear models (GLMs) used for sgRNA-level CRISPR-KO screen analyses for NK cell tumor infiltration in four tumor models. Coefficients are only shown for the top 50 highly variable genes.

**Extended Data Figure 3:**
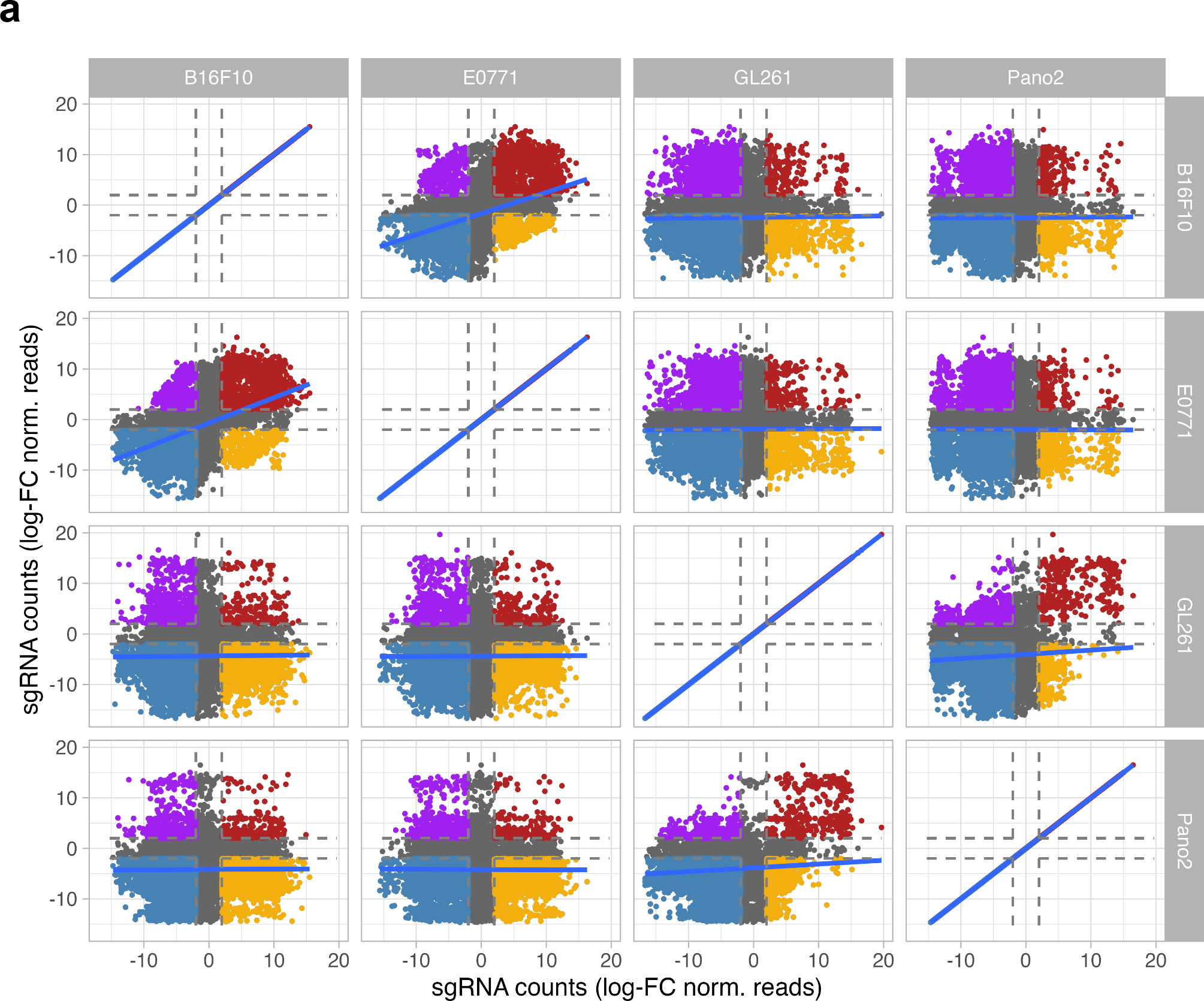
Comparisons between independent, parallel in vivo CRISPR-KO screens for NK cell tumor infiltration. **a,** Scatter plots that compare the sgRNA log fold-changes for the tumor infiltration screen readouts in four independent CRISPR-KO screen tumor models.

**Extended Data Figure 4:**
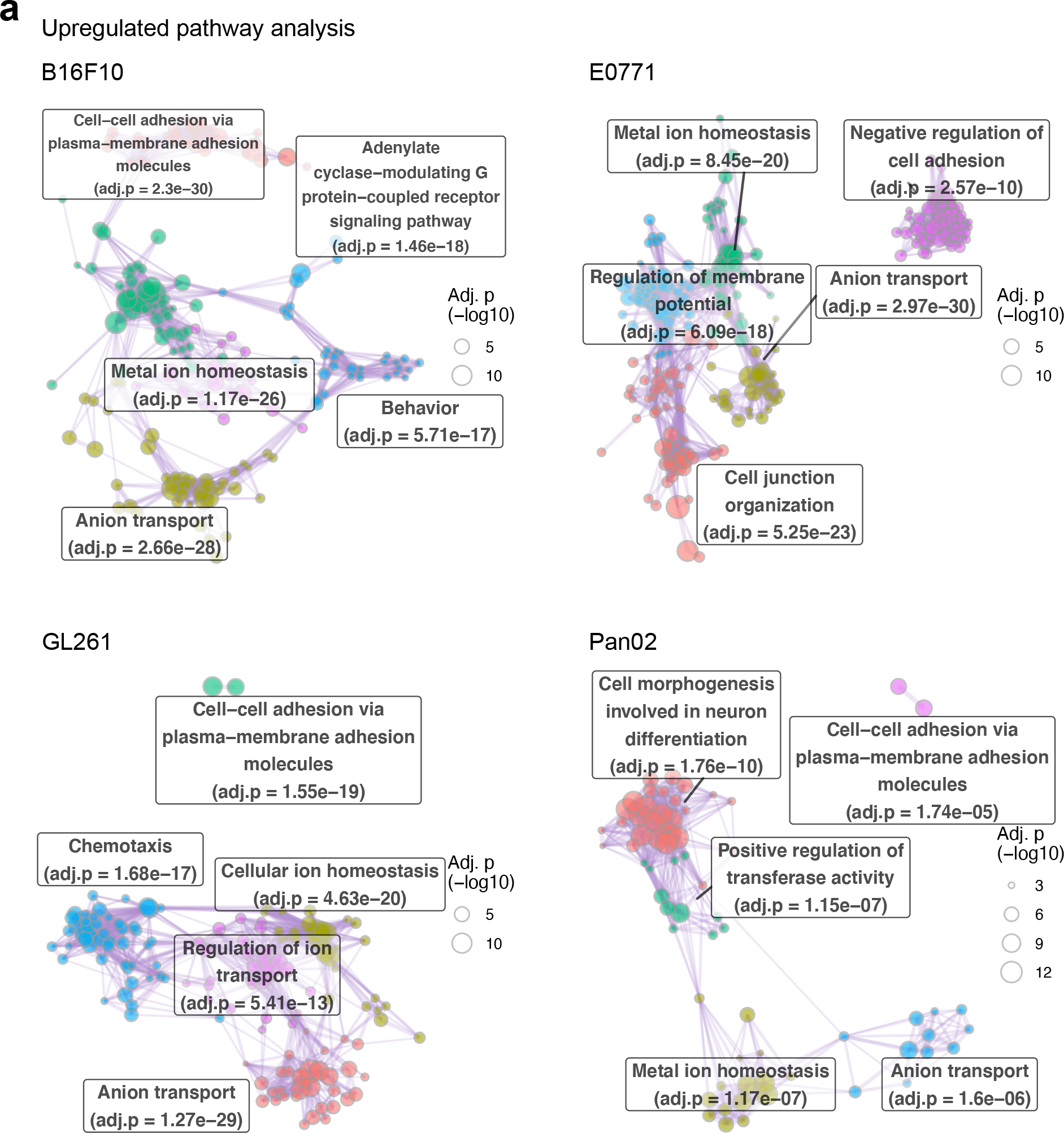
Meta-pathway analyses of parallel in vivo CRISPR-KO screens for NK cell tumor infiltration. **a,** Network plots of clustered gene ontology terms from pathway analyses of upregulated genes in the in vivo CRISPR-KO screens for NK cell tumor infiltration performed in four independent tumor models. Pathway enrichment analyses were performed by gProfiler2, and significantly enriched pathways were clustered with Leiden algorithm. Pathway clusters are labeled by meta-pathway of that cluster (see methods for details). The top five meta-pathways are shown for each plot.

**Extended Data Figure 5:**
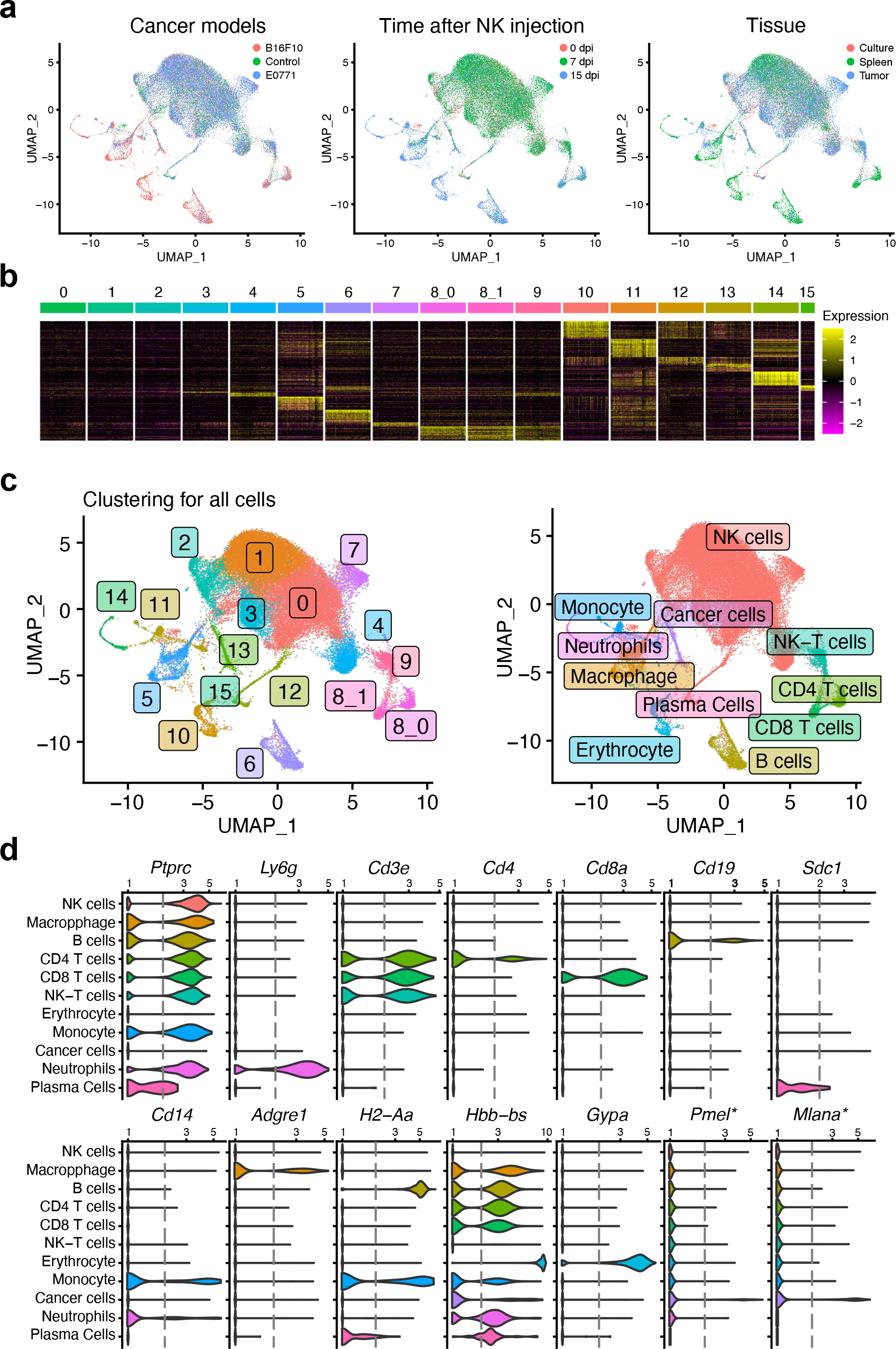
Single-cell transcriptomic analysis of cancer-immune cell populations across time in different tumor models and tissues. **a,** UMAP plots of cells across nine integrated single-cell transcriptomic datasets. Cells are color-labeled according to their original dataset, including pre-transfer donor NKs (controls) and cells from different timepoints, tumor models, and tissues. **b,** Heatmap of discreet expression patterns across different cell population subsets. Scaled gene expression is presented for 100 representative cells of each population. **c,** UMAP plot of cell subset populations with unlabeled and labeled cells in the left and right panels, respectively. **d,** Violin plots of the expression of select immune marker genes, compared across different cell subset populations. The distribution of log-scaled expression data is shown for each cell subset, and a dashed line represents a log-scale expression threshold of 2.

**Extended Data Figure 6:**
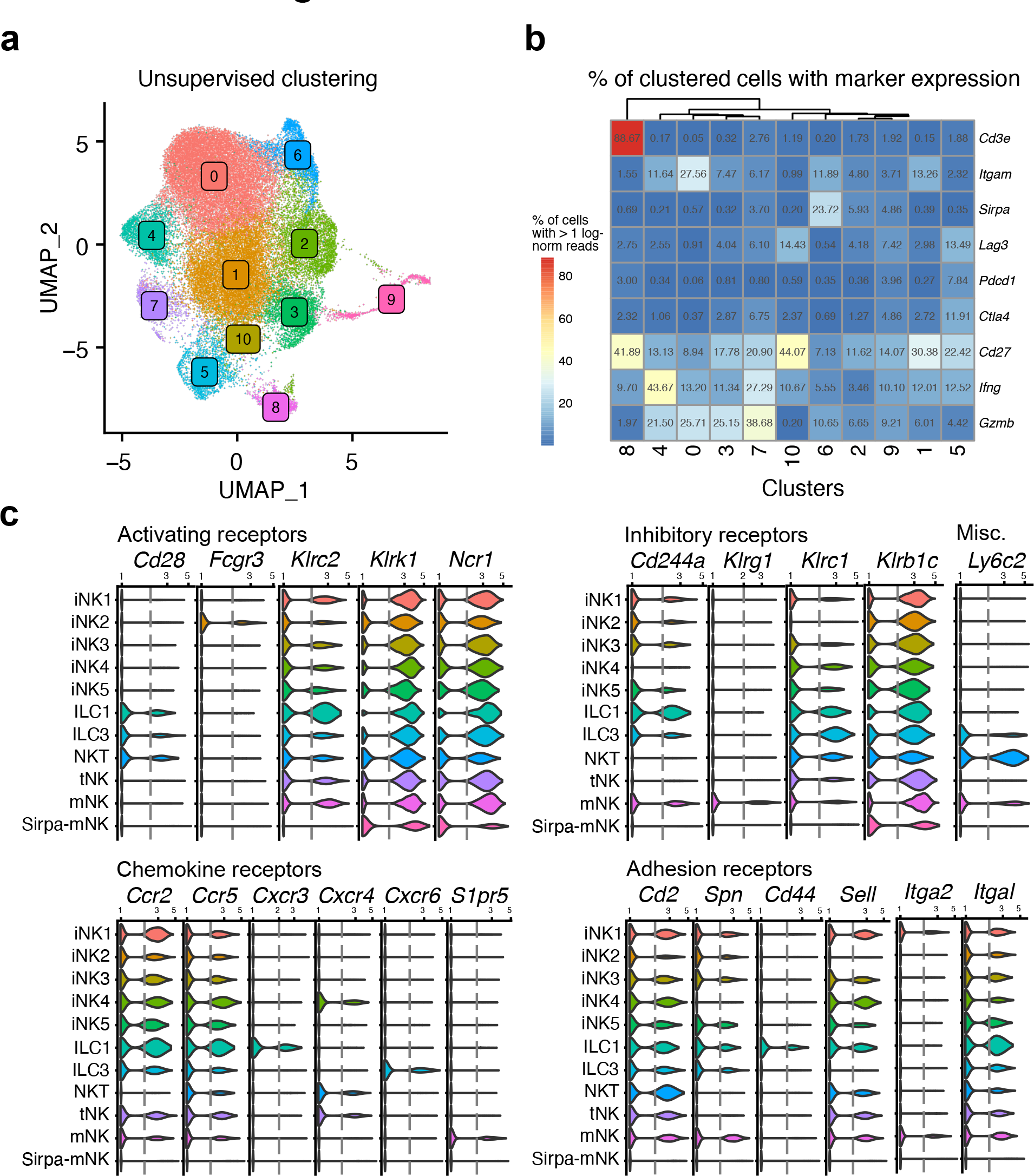
Single-cell transcriptomic analysis of NK cell populations across time in different tumor models and tissues. **a,** UMAP plot of unlabeled cell subset populations determined from unsupervised clustering. **b,** Heatmap of the percent of cells from each cluster that expresses select immune marker genes at > 1 log-normalized reads. **c,** Violin plots of the expression of select immune marker genes, compared across different cell subset populations. The distribution of log-scaled expression data is shown for each cell subset, and a dashed line represents a log-scale expression threshold of 2.

**Extended Data Figure 7:**
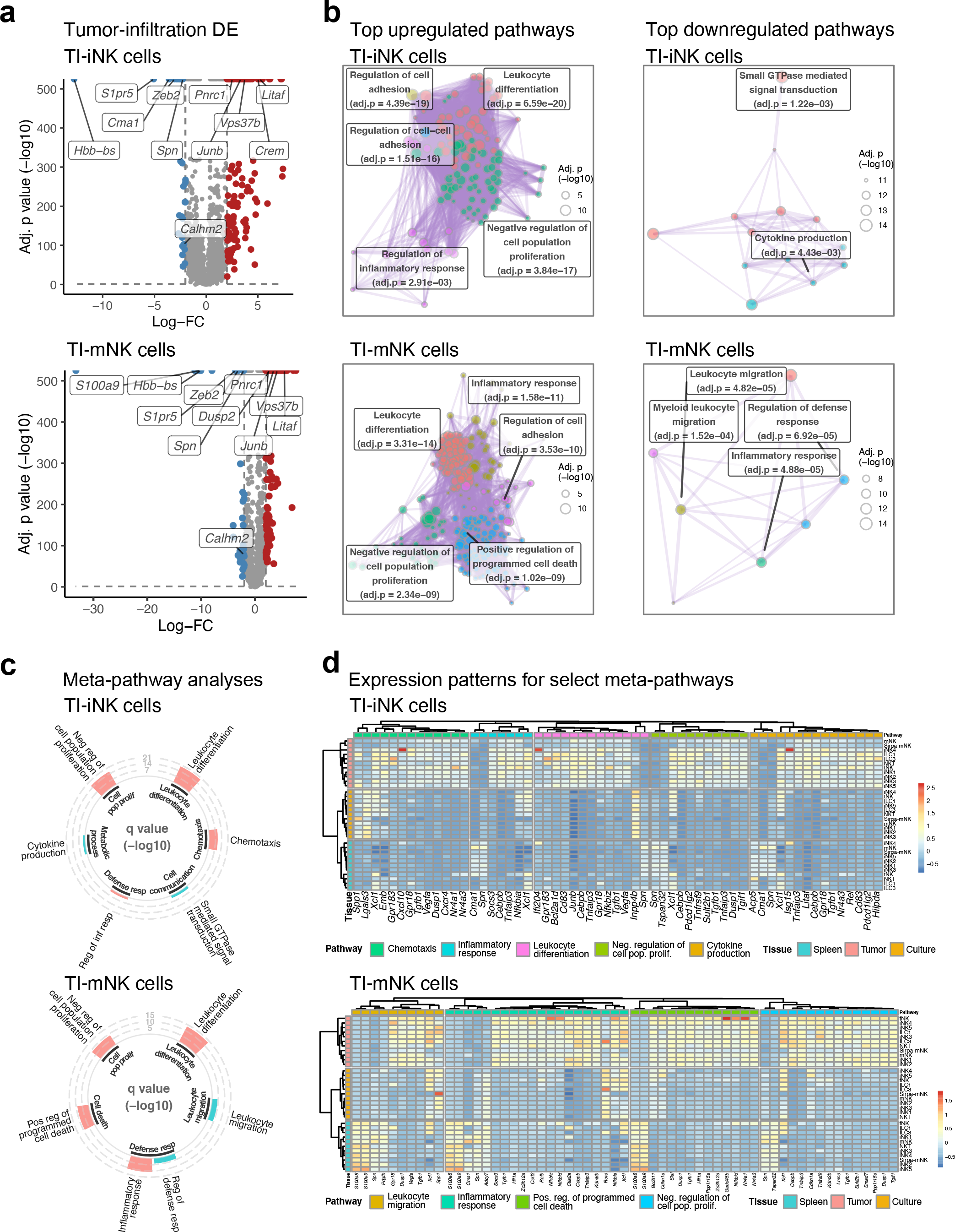
Differential expression analyses of tumor infiltration in immature and mature NK single-cell data. **a,** Volcano plots of differential expression (DE) analyses for tumor infiltration (TI) of immature (iNK) and mature NK (mNK) single-cell data (top and bottom panels, respectively). DE analyses were performed using single-cell expression data fitted to Gamma-Poisson generalized linear models (shown above plots), and quasi-likelihood F tests. Upregulated and downregulated genes are shown by respective red and blue dots (q < 0.01, absolute log-FC > 2), and the top DE gene names are presented for each. **b,** Network plots of clustered gene ontology terms from pathway analyses of upregulated genes or downregulated genes (left and right panels, respectively) in the DE analysis of TI in iNK and mNK single-cell expression data (top and bottom panels, respectively). Pathway enrichment analyses were performed by gProfiler2, and significantly enriched pathways were clustered with Leiden algorithm. Pathway clusters are labeled by meta-pathway of that cluster (see methods for details). The top five meta-pathways are shown for each plot. **c,** Circular bar plots of meta-pathway analysis results for DE genes related to TI in iNK and mNK single-cell expression data (top and bottom panels, respectively). Meta-pathways are shown for relevant immune-related categories, and pathway significance is represented by bar height. Meta- pathways from the analyses of upregulated and downregulated DE genes are indicated by red and blue bars, respectively. **d,** Heatmaps of select meta-pathways from the enrichment analyses of TI in iNK and mNK single-cell expression data (top and bottom panels, respectively). For each DE gene, population-averaged scaled-normalized gene expression is presented by color. Rows and columns are color-annotated by NK subset/tissue and meta-pathway, respectively. Hierarchical clustering was separately performed within each meta-pathway using highly variable DE genes (> 1 sd), and duplicate DE genes were allowed across meta-pathways.

**Extended Data Figure 8:**
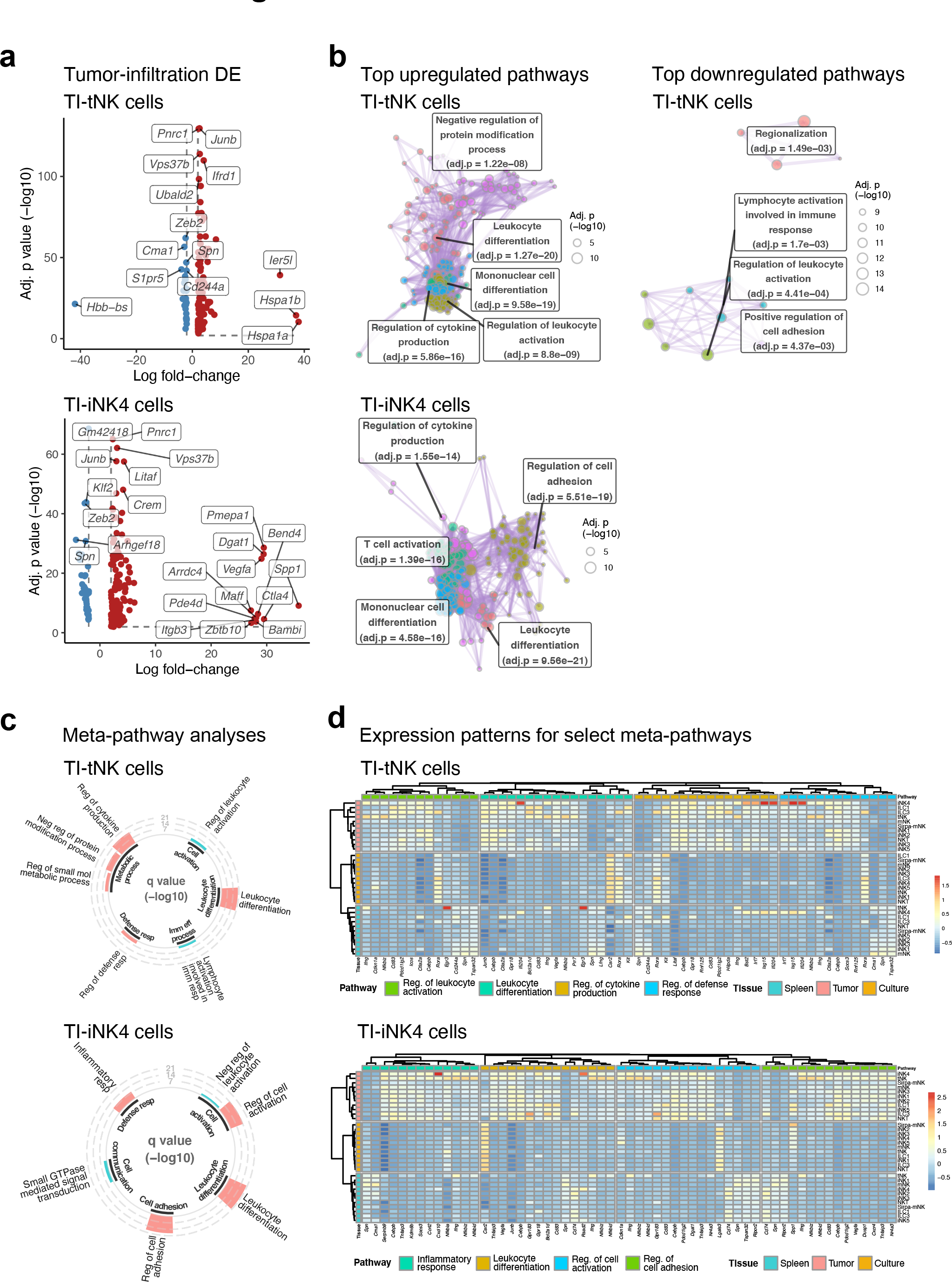
Differential expression analyses of tumor infiltration in transitional NK and the immature NK 4 subset single-cell data. **a,** Volcano plots of differential expression (DE) analyses for tumor infiltration (TI) of transitional (tNK) and immature NK population 4 (iNK4) single-cell data (top and bottom panels, respectively). DE analyses were performed using single-cell expression data fitted to Gamma-Poisson generalized linear models (shown above plots), and quasi-likelihood F tests. Upregulated and downregulated genes are shown by respective red and blue dots (q < 0.01, absolute log-FC > 2), and the top DE gene names are presented for each. **b,** Network plots of clustered gene ontology terms from pathway analyses of upregulated genes or downregulated genes (left and right panels, respectively) in the DE analysis of TI in tNK and iNK4 single-cell expression data (top and bottom panels, respectively). Pathway enrichment analyses were performed by gProfiler2, and significantly enriched pathways were clustered with Leiden algorithm. Pathway clusters are labeled by meta-pathway of that cluster (see methods for details). The top five meta-pathways are shown for each plot. **c,** Circular bar plots of meta-pathway analysis results for DE genes related to TI in tNK and iNK4 single-cell expression data (top and bottom panels, respectively). Meta-pathways are shown for relevant immune-related categories, and pathway significance is represented by bar height. Meta- pathways from the analyses of upregulated and downregulated DE genes are indicated by red and blue bars, respectively. **d,** Heatmaps of select meta-pathways from the enrichment analyses of TI in tNK and iNK4 single-cell expression data (top and bottom panels, respectively). For each DE gene, population-averaged scaled-normalized gene expression is presented by color. Rows and columns are color-annotated by NK subset/tissue and meta-pathway, respectively. Hierarchical clustering was separately performed within each meta-pathway using highly variable DE genes (> 1 sd), and duplicate DE genes were allowed across meta-pathways.

**Extended Data Figure 9:**
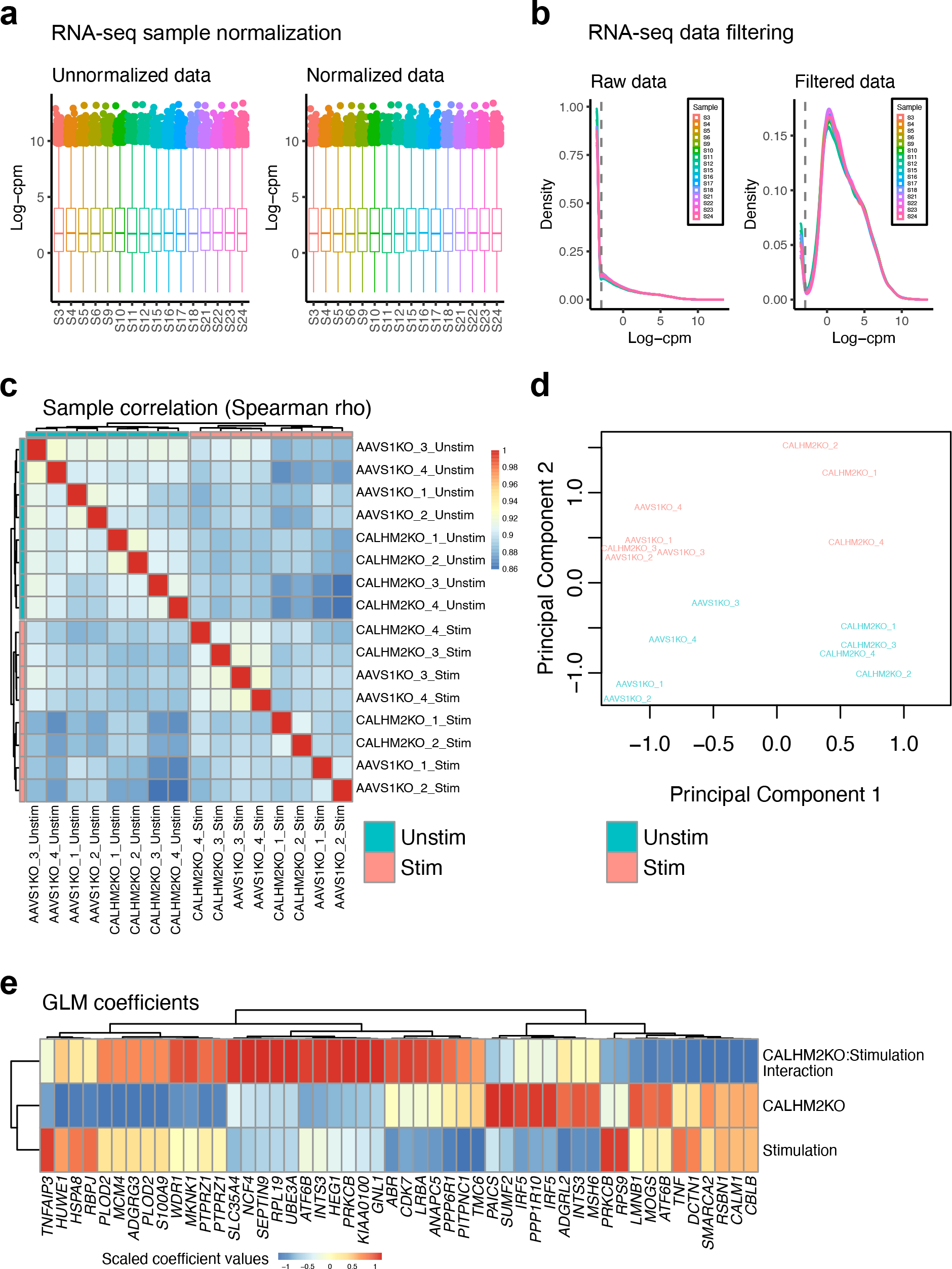
Quality control metrics for RNA-seq analysis of CALHM2-KO in α-HER2 CAR-NK cell samples. **a,** Box-whisker plot of unnormalized and normalized read count distributions across all α-HER2 CAR-NK cell samples. Distributions are presented with log2 counts per million reads (CPM) for each sample. **b,** Density plots of raw and filtered RNA-seq data (log2 CPM) for all α-HER2 CAR-NK cell samples. **c,** Heatmap of the Spearman correlation between CAR-NK cell samples included in the bulk RNA-seq data analysis. **d,** Scatter plots for the multidimensional scaling of processed RNA-seq data of CAR-NK cell samples. **d,** Heatmap of key experimental coefficients in the generalized log-linear model (GLM) of the RNA-seq analysis of CALHM2-KO in CAR-NK cells with and without stimulation. Coefficients are only shown for the top 50 highly variable genes.

**Extended Data Figure 10:**
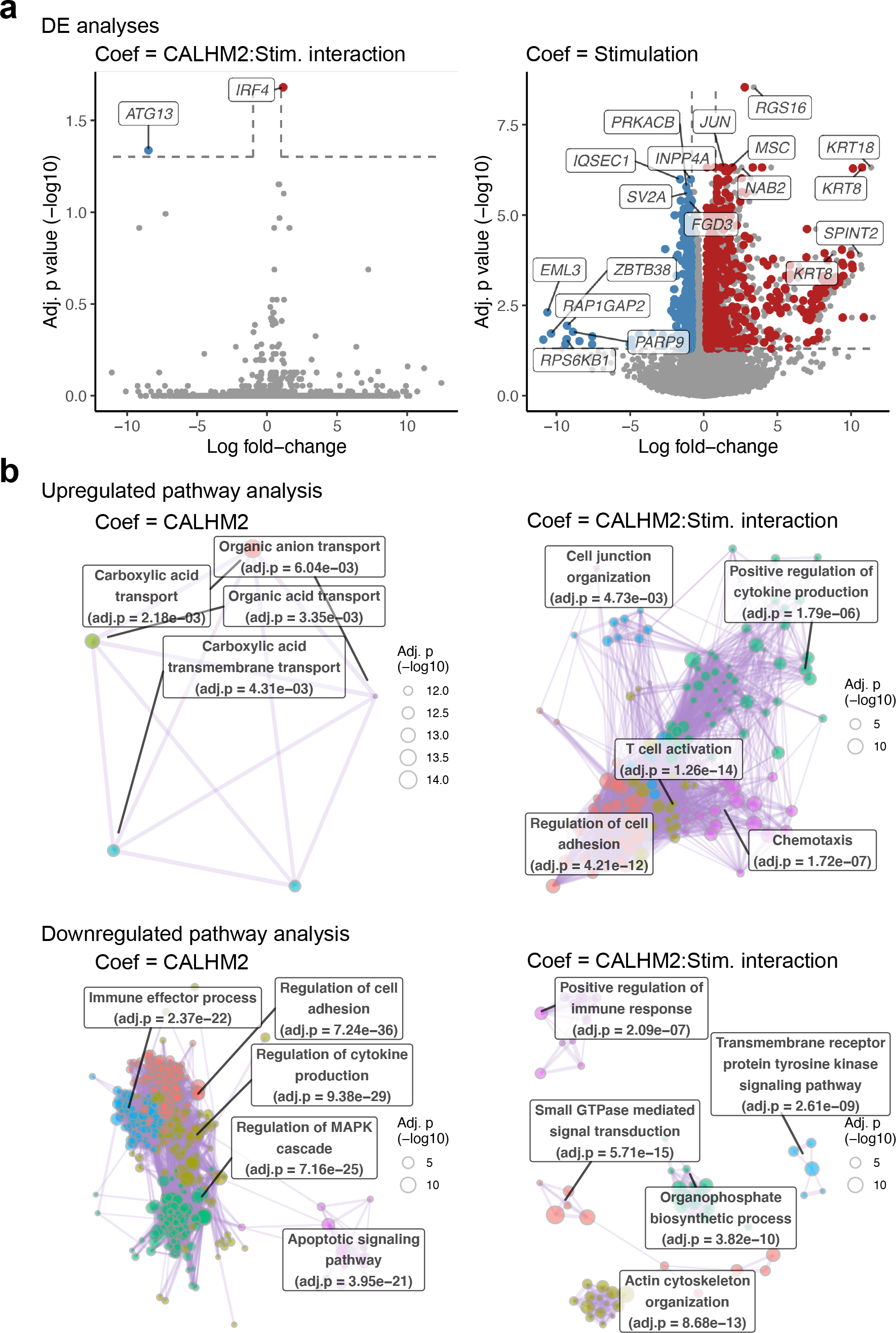
Differential expression analyses of CALHM2-KO in α-HER2 CAR-NK cells. **a,** Volcano plots of the differential expression (DE) analysis of CALHM2:stimulation interaction and stimulation in human donor α-HER2 CAR-NK cells. DE analysis was performed using bulk RNA-seq expression data fitted to a negative binomial generalized linear model that included coefficients for donor paired samples (CALHM2/AAVS1 gRNA), stimulation status, and the interaction between CALHM2-KO and stimulation. The effect of CALHM2-KO was assessed by quasi-likelihood F tests. Upregulated and downregulated transcripts are shown by respective red and blue dots (q < 0.05, absolute log-FC > 0.8), and the top significant DE gene names are labeled. **b,** Network plots of clustered gene ontology terms from pathway analyses of upregulated genes or downregulated genes (top and bottom panels, respectively) in the DE analysis of CALHM2-KO or the interaction of CALHM2-KO:Stimulation (left and right panels, respectively). Pathway enrichment analyses were performed by gProfiler2, and significantly enriched pathways were clustered with Leiden algorithm. Pathway clusters are labeled by meta-pathway of that cluster (see methods for details). The top five meta-pathways are shown for each plot.

**Extended Data Figure 11:**
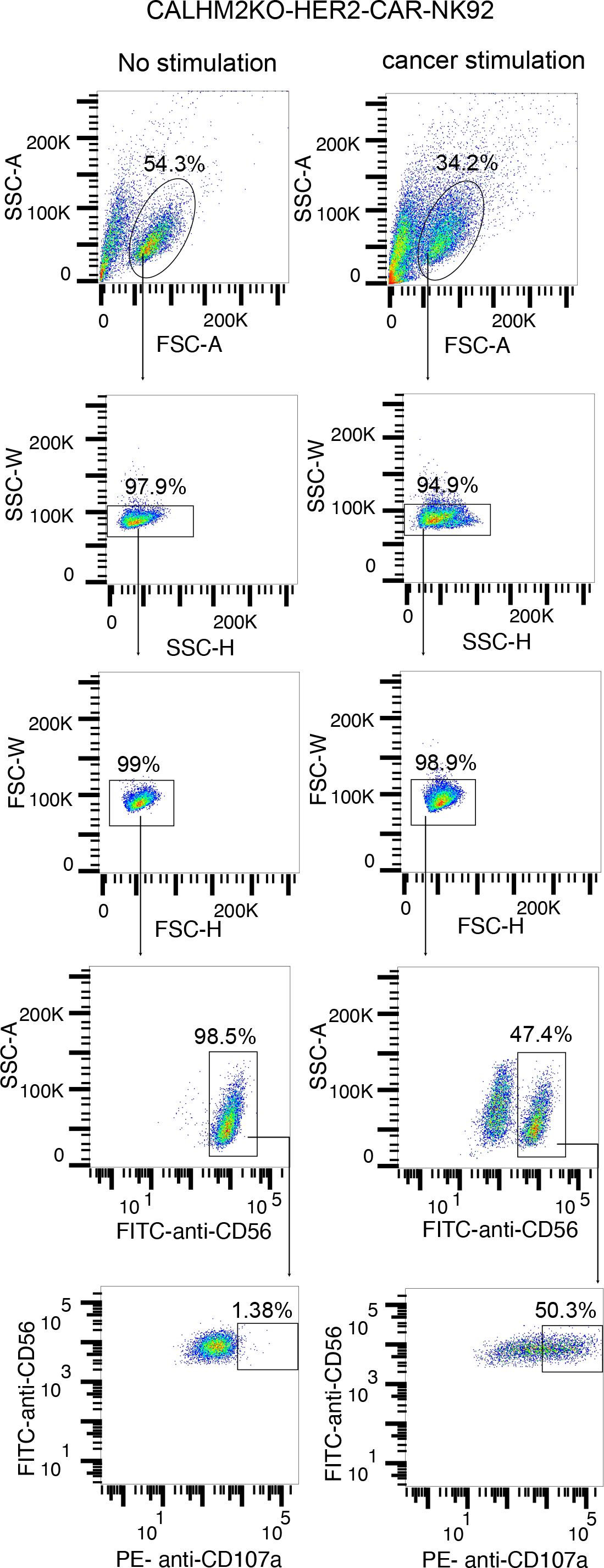
Representative flow gating plot.

**Supplementary Table:** Reagent and resource information.

Non-NGS Source Data

Non-NGS source data and statistics provided in an excel file: NK screen_source data_V1b.xlsx

NGS Source Data

NGS source data and statistics provided in zip files:

**Source Data 1:** Screen analysis source data.

**Source Data 2:** Screen meta-pathway analysis source data.

**Source Data 3:** Single-cell RNA-seq population analysis source data.

**Source Data 4:** Single-cell RNA-seq DE analysis source data.

**Source Data 5:** Single-cell RNA-seq meta-pathway analysis source data.

**Source Data 6:** Bulk RNA-seq DE analysis source data.

**Source Data 7:** Bulk RNA-seq meta-pathway analysis source data.

